# Relative Risk Assessment of Ecological Areas with the Highest Potential Impact of Underwater Released Exhaust CO_2_ from Innovative Ships

**DOI:** 10.1101/2021.09.30.462553

**Authors:** Yuzhu Wei, Michel Kroeze, Csilla Vámos, Edwin M. Foekema, Ron van Lammeren, Albertinka J. Murk

**Affiliations:** Marine Animal Ecology group, Wageningen University and Research, P.O. box 338, 6700 AH Wageningen, The Netherlands; Wageningen Marine Research, Wageningen University and Research, P.O. Box 57, 1780 AB Den Helder, The Netherlands; Laboratory of Geo-Information Science and Remote Sensing, Wageningen University and Research, P.O. Box 8130, 6700 EW Wageningen, The Netherlands

## Abstract

Applying underwater released exhaust gas as ‘air lubrication’ along the ship’s hull to reduce the energy consumption is under development. However, this direct emission to the water could pose a risk to the local marine environment, especially in shipping-dense areas. Specifically, CO_2_, a dominant component in the exhaust gas, has the potency to enhance algal blooms and cause acidification. This study provides the first relative risk assessment of ships with underwater release exhaust gas systems on a global scale, taking into account local water conditions and shipping intensity. Risk was characterized for 262 marine ecoregions by plotting the expected CO_2_ emission from ships to water against the estimated vulnerability to acidification and algal blooms. The vulnerability of each ecoregion was assessed based on background dissolved inorganic carbon (DIC) level, chlorophyll-a concentrations and total alkalinity. The results reveal that areas with relatively high vulnerability are mainly located above 30° N latitude. The Yellow Sea, Southern China Sea, and North Sea come out as relatively high risk areas. Looking in more detail to European high-risk ecoregions, the highest risk levels are found in areas with dense shipping lanes and maritime chokepoints, e.g. the Strait of Dover and the Strait of Gibraltar. This was the first attempt to make such a risk assessment and the outcome is only indicative. In a next phase additional parameters, such as water currents and biological composition of the ecosystem should be included.

**Key Points:**

- The first global environmental risk assessment was made for innovative ships that apply underwater released exhaust gas as ‘air lubrication’
- Ecoregions with relatively high vulnerability to acidification and algal blooms are mainly located above 30° N latitude
- The risk for the marine ecosystem of the underwater released exhaust CO_2_ from ships is limited to specific shipping-dense areas

## 1 Introduction

Underwater released exhaust gas systems as ‘air-lubrication’ of ship hulls are under development and expected to reduce the drag force, in turn reducing the fuel consumption and total emission of global shipping activities (Sapra et al., 2017; Van Biert et al., 2016). Additionally, it improves the air quality on working decks (Sapra et al., 2017). These advantages make the system attractive, especially under the more strict maritime emission regulation (Sapra et al., 2017; Van Biert et al., 2016), e.g. the nitrogen oxides (NO_x_) emissions control (IMO, 2013). However, such an application implies concentrated input of exhaust gas in the local marine ecosystem and may pose a serious risk, especially in the case of intensively used shipping lanes. Specifically, CO_2_, one of the main components in the exhaust gas (Anderson et al., 2015), could enhance algal bloom and cause acidification, depending on the local environmental conditions.

For photosynthetic organisms, dissolved CO_2_ is one of the essential nutrients (together with N, P, Fe, etc.). CO_2_ is continuously exchanged between the atmosphere and seawater phases, which provides carbon nutrients to these aquatic primary producers, like microalgae (Markou et al., 2014; Zeebe & Wolf-Gladrow, 2001). In the case of high algal densities, the mass transfer of CO_2_ from the atmosphere to the liquid state can be slower than the algal uptake rate (Markou et al., 2014; Zeebe & Wolf-Gladrow, 2001). As long as other nutrients are still available, CO_2_ availability will then become the factor determining further algal growth (Markou et al., 2014). In such conditions, underwater released exhaust CO_2_ may relieve this carbon resource limitation and stimulate further development of algae densities potentially resulting in an algal bloom or extending the blooming period.

After release, CO_2_ dissolves in water and increases the Dissolved Inorganic Carbon (DIC) level (Zeebe & Wolf-Gladrow, 2001). With increasing DIC levels, the seawater becomes acidified, as indicated by a reduction of the pH value. This acidification may e.g. inhibit the calcification process of organisms such as corals and mussels (Kurihara, 2008; Sunday et al., 2017). The inhibition of the calcification process hampers the organism’s ability to form calcium carbonate (CaCO_3_) structures like skeletons and shells (Fassbender et al., 2016) with negative impact on growth and survival (Kurihara, 2008; Wei et al., 2019). The capacity of seawater to resist a decrease in pH with increasing CO_2_ concentrations mainly depends on the amount of anions present (e.g. CO_3_^2-^ and HCO_3_^-^) (Zeebe & Wolf-Gladrow, 2001) and is indicated by Total Alkalinity (TAlk). Waters with a higher TAlk have stronger acid-neutralizing capacity and can absorb larger amounts of CO_2_ before the water pH drops (Omernik & Powers, 1983). The sensitivity of marine regions for enhanced CO_2_ exposure therefore will depend on the local TAlk level and nutrient levels, information that can be derived from data described for global marine ecoregions (Spalding, Fox et al. 2007) and can be used in ArcGIS to combine different conditions.

Both, stimulation of the algae density growth and inhibition of the growth of calcifying organisms, can be devastating for local marine ecosystems. Of course, the intensity of the CO_2_ exposure as well as the sensitivity of the receiving marine ecosystem will determine the eventual risk for adverse effects to develop. The present study aims to assess for ecoregions the relative risk that projected underwater released exhaust CO_2_ will cause algal blooms or acidification.

## 2 Materials and Methods

To be able to assess the risk of underwater exhausted CO_2_ for the local marine environment, both the exposure, as well as the vulnerability to CO_2_ needed to be quantified and compared for each ecoregion (Figure 1) (section 2.4). The exposure level was assessed from the reported CO_2_ emission from maritime traffic in 2018 in combination with the assumed saturation level of DIC in seawater (section 2.2). Quantification of the vulnerability to CO_2_ was based on two indicators: 1) TAlk level as a measure of resistance to acidification, and 2) chlorophyll-a concentration as a measure of eutrophication (section 2.3).

### 2.1 Data collection, representation and projection

All pre-processing and spatial analyses were performed in ESRI ArcGIS 10.6.1 unless described otherwise. All datasets were converted to the Compact Miller projection prior to further analyses.

#### 2.1.1 Global ocean map data collection

The coastlines of the global map were created using the land-sea mask dataset and methods as described by Halpern et al. (2008). The land-based data that in reference to this data mask occurred within the ocean, or ocean based data that occurred on land were clipped and removed to ensure consistency across all data used in our analysis (Halpern et al., 2008). All data were represented at 1 km^2^ resolution. For the collected data that was only available at a coarser native resolution, it was assumed that the coarse-scale value was evenly distributed across all 1 km^2^ cells within that region (Halpern et al., 2015). This essentially maintains the coarse-scale pattern while the finer resolution information is preserved when it is available. For the gaps in datasets, the null values were filled with a 5 x 5 focal mean filter (Sharma et al., 2010; Tomlin, 2016). It was specified that only cells containing values were used to find the focal mean of the target cell. This means that a null value will be ignored when it exists within the neighbourhood of the focal mean filter. Thus, this process removed gaps while minimizing unjustified smoothing effects of missing data on the dataset (Sharma et al., 2010; Tomlin, 2016). The filter was run multiple times from the edge of each gap until all encapsulated null values were filled (i.e. null values surrounded by values). Very large data gaps occurred in areas with sea ice. These data gaps were not filled due to their large size. Instead, ice masks were created to indicate where scores are less certain. Ice masks were created to indicate where scores are less certain. To indicate which areas are counted as sea ice, the daily fractional ice cover data from 2012 were obtained via the Advanced Very High Resolution Radiometer (AVHRR) Pathfinder Version 5.2 (Casey et al., 2011). The daily fractional ice cover data were then averaged by meteorological season and converted to binary terms: either the cell has ice, or the cell has no ice. Grid cells that contained an average sea ice fraction greater than 0.15 were considered to have ice, and were thus included in the ice masks.

**Figure 1.**
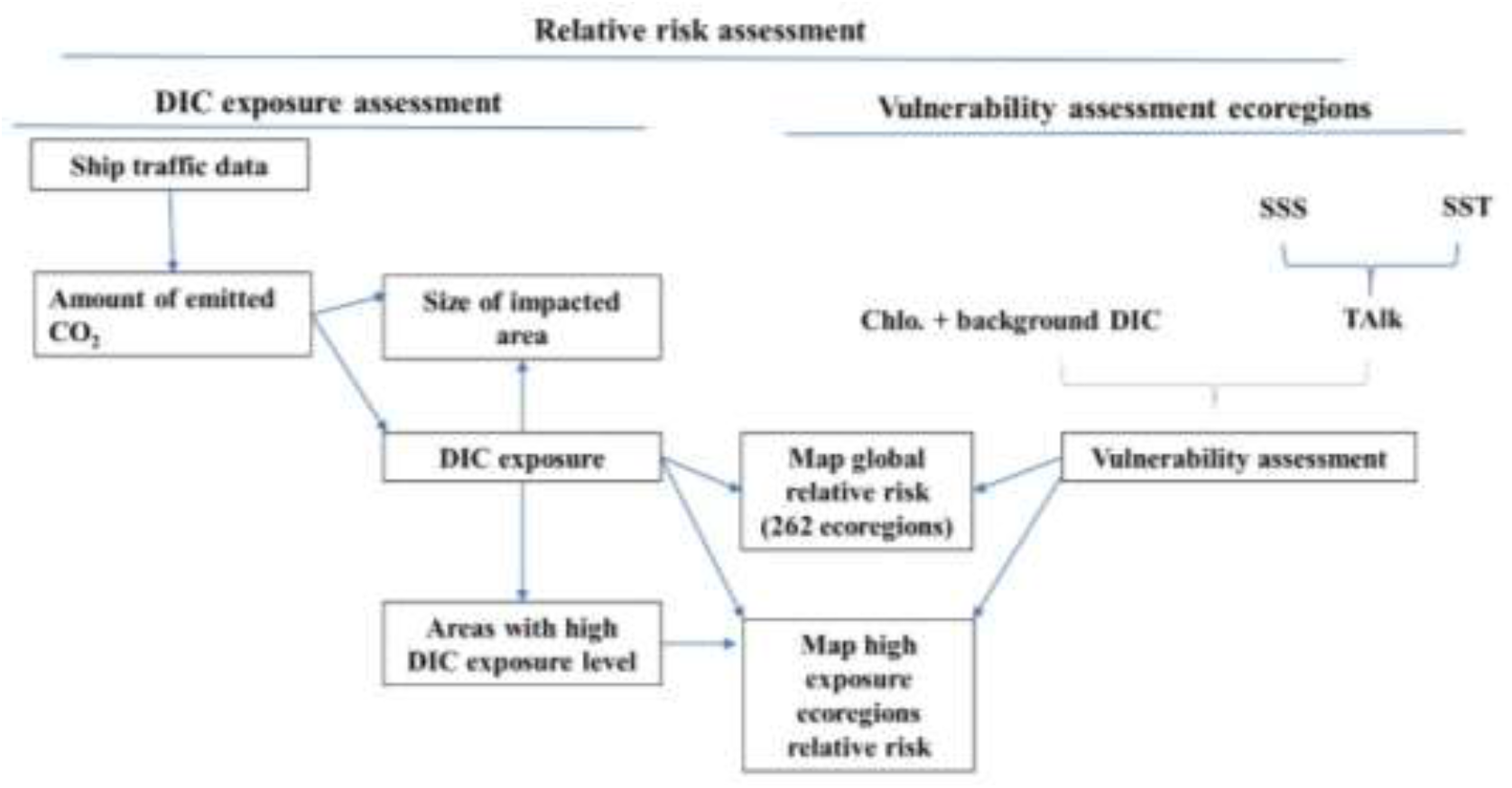
Workflow diagram of the relative risk assessment of underwater released exhaust CO_2_ from maritime shipping for global and local ecoregions. Dissolved inorganic carbon (DIC) exposure assessment of CO_2_ emitted from maritime shipping and vulnerability assessment of algal bloom (indicated as Chlo. + original DIC level) and acidification (indicated as TAlk level) were considered in risk characterization for global and local ecoregions to CO_2_ emitted by underwater exhausts. SSS: seawater surface salinity; SST: seawater surface temperature; Chlo.: chlorophyll-a; TAlk: Total Alkalinity.

#### 2.1.2 Ecoregions

The global ocean was divided into 262 ecoregions by taking two steps: 1) the coastal and shelf waters shallower than 200m were divided into 232 ecoregions based on biogeographic patterns (Mark D Spalding et al., 2007a), and 2) all remaining open oceans were divided into 30 ecoregions following similar classifications as the coastal and shelf water areas (Mark D Spalding et al., 2012). Each ecoregion consists of relatively homogeneous species composition that is distinct from adjacent areas (Mark D Spalding et al., 2007a). The datasets were collected via the Nature Conservancy’s Geospatial Conservation Atlas (Atlas, 2019) and UNEP’s Ocean data viewer (Conservancy, 2012).

As Europe was later indicated to be continent that experiences the highest DIC exposure, the 15 marine ecoregions around Europe (Figure S1a in the supporting information) were subdivided into ecologically relevant classes based on ocean floor depth (Waller, 1996): the epipelagic zone (0 – 200 m), the mesopelagic zone (200 – 1000 m), the bathypelagic zone (1000 – 2250 m), the abyssopelagic zone (2250 – 4500 m), and the hadopelagic zone (4500 – 11500 m). To show variation in risk level between coastlines and open seas, the epipelagic zone was arbitrarily sub-divided into five depth classes, resulting in a total of 9 depth classes (Figure S1b in the supporting information). Dividing the 15 European ecoregions by depth class resulted in a total of 114 European sub-ecoregions that were labelled according to the ecoregion code (same as the global ocean code) and depth class.

#### 2.1.3 Chlorophyll-a concentration and DIC

Nutrient concentrations in seawater are regularly measured on a local scale, but a dataset with global coverage that met our needs was not available. As an alternative, the chlorophyll-a concentration was used as an indicator for algal biomass, since the chance that additional DIC will promote further algal development is greater at higher algal density. The chlorophyll-a dataset was collected from NASA’s Aqua-MODIS satellite (NASA Goddard Space Flight Center, 2018) for each calendar month of 2018 and aggregated to seasonal means (December - February, March - May, June - August, and September - November). The yearly average DIC concentrations of the ocean were collected from the GLODAPv2 dataset (Lauvset et al., 2016; Olsen et al., 2016).

#### 2.1.4 TAlk

As the vulnerability indicator for acidification, TAlk was calculated based on sea surface temperature (SST) and sea surface salinity (SSS) by following the equation described by Lee et al. (2006). The SSS and SST data for each calendar month of 2018 were collected from NASA’s JPL SMAP satellite (Nasa/Jpl, 2019) and NOAA’s Coral Reef Watch satellites (Watch, 2018, updated daily) respectively.

### 2.2 DIC exposure assessment

#### 2.1.1 Maritime CO2 emission

To assess the DIC exposure level caused by underwater released exhaust CO_2_, the amount of maritime emitted CO_2_ has to be estimated. For this, a 2013 shipping intensity dataset was collected from the Knowledge Network for Biocomplexity (Halpern et al., 2015). This dataset contains the registered number of ships in each 1 km^2^ grid cell (X) in a log[*X* + 1] transformed and rescaled (between 0 and 1, with the highest per-pixel transformed value = 1) form. The emission factors for each ship class were collected from the literature (ECTA, 2011; McKinnon & Piecyk, 2010; Otten et al., 2017).

#### 2.1.2 Shipping intensity and the amount of emitted CO2

To calculate the number of ships in each grid cell (X), the rescaling of the collected shipping intensity dataset (occurring in a rescaled log[*X* + 1] form) was reversed first using the following formula:

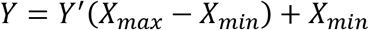

Here *Y′* is the log [X+1] transformed data, *Y* is the collected data and X_min_ and X_max_ are the minimum and the maximum number of registered ships in each grid cell. The collected dataset contains zero, which is the result of “log 1”. Thus, X_min_ is zero. The unknown X_max_ was estimated by log[*X* + 1] transforming the raw ship traffic dataset from Halpern et al. (2015)’s study. This raw dataset contains the number of ships in each grid cell, but also includes invalid and cross land routes. Finally, the number of registered ships in a grid cell (X) of 2013 was calculated by the exponential of the log transformed data (2).

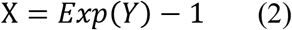

Next, the calculated data were multiplied with the 8.16% growth rate (from 2013 to 2018) in the global merchant fleet to project the ship traffic intensity for 2018 (UNCTAD, 2018a). The presented study focused on maritime shipping in 2018, because that was the most recent and reliable maritime shipping information available from the United Nations annual review of maritime transportation, when this study was carried out. This adaption also created a worst-case-scenario as it assumes that all vessels are merchant ships.

Finally, the total amount of emitted CO_2_ per ship per grid cell (E_T_) is quantified by multiplying the proportional share (*P_i_*) of each *i… m* vessel class, the amount of emitted CO_2_ per km (*K_i_*) of each *i…m* vessel class, and the average travel distance of a ship to pass a 1 km^2^ grid cell: 0.7 km (Text S1 in the supporting information) (3). To consider the worst-case-scenario, *K_i_* was determined based on the highest emission factors in Table 1. An emission factor of the ‘other ships’ class was not available because of the broad definition of this class (including all liquefied petroleum gas tankers, ferries, cruises, etc.) by UNCTAD (2018b). Therefore, the emission factor of ‘other ships’ was assumed to be in the same range as ‘general cargo’, since both classes share a similar average deadweight tonnage (dwt) and short-sea shipping function (Table 1).

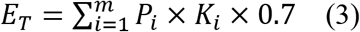

**Table 1.** Information of Glbal Maritime Merchant in 2018.

| Merchant ship class | Nr. ship | Mean dwt/ship (tonnes) | Emission factors (lower and upper levels) (g CO<sub>2</sub> /ton/km<sup>a</sup>) | kg CO<sub>2</sub> emitted/km/ship<sup>d</sup> |
| --- | --- | --- | --- | --- |
| Oil tankers | 10420 | 53839 | 10.3 - 15 | 807.6 |
| Bulk carriers | 11125 | 73618 | 7 - 11.9 | 876.1 |
| Container ships | 5164 | 48993 | 8.4 - 12<sup>b</sup> | 587.9 |
| General cargo | 19613 | 3773 | 13.9 - 21 | 79.2 |
| Other vessels | 47847 | 4535 | 13.9 - 21<sup>c</sup> | 95.2 |

*Note.* The number of global maritime merchant ships in each purpose class (nr. ship), the mean deadweight tonnage (dwt) of each ship in 2018 (UNCTAD, 2018b), the corresponding CO<sub>2</sub> emission factors (ECTA, 2011; McKinnon & Piecyk, 2010; Otten et al., 2017), as well as the mean kg CO<sub>2</sub> emitted per traveled km per ship. <sup>a</sup>Unit defining transport performance. An output of one is reached when one ton is transported one km. <sup>b</sup>Emission factors of container ships based on twenty-foot equivalent units (TEU's) instead of dwt. <sup>c</sup>Emission factors are assumed to be the same as for ‘general cargo’ due to their similar mean dwt and short-sea shipping function. <sup>d</sup>Calculated as: (mean dwt per ship x emission factor) / 1000.

#### 2.2.3 Emitted CO2 induced DIC exposure

For the risk assessment, we assumed that all CO_2_ produced by the ships is emitted underwater. The water DIC level is determined by the amount of dissolved CO_2_ instead of the total emitted CO_2_ (Figure 1). Potentially, all CO_2_ can dissolve in water and change the carbon composition in water. However, when the saturation of DIC is reached (depending on e.g. temperature, salinity and pressure), additional CO_2_ will not dissolve in the water anymore (Zeebe & Wolf-Gladrow, 2001) and will escape to the atmosphere. In addition, the dissolved CO_2_ continually exchanges with the CO_2_ gas in the atmosphere, which eventually leads to an equilibrium. Due to the saturation of DIC in water and the equilibrium of CO_2_ concentrations between water and atmosphere, increasing the amount of injected CO_2_ in DIC saturated water does not raise the exposure of the local marine ecosystem further. Therefore, the saturation level of DIC and the equilibrium of CO_2_ concentrations in water and atmosphere phases determine the maximum exposure level caused by underwater released exhaust CO_2_.

#### 2.2.4 Determination of the DIC saturation level

We determined the maximum saturation level of DIC in seawater and the time required to reach this in a laboratory experiment. For this, a flow of air (flow rate 148 ml/min) with 5% CO_2_ was continuously injected into 150ml of artificial seawater (sea salt Marine Zoomix®; temperature 20 °C; salinity 31.7‰; alkalinity 2.07 mmol/l and background pH 8.1). The uptake of CO_2_ in the water was reflected by the lowering of the pH. The injection was terminated when the water pH level did not further decrease indicating that the maximum saturation was reached. After terminating the CO_2_ supply, the water pH level started increasing as the surplus CO_2_ from the water escaped to the atmosphere. The water pH was continually measured until the pH was stable, and the equilibrium of CO_2_ in the water and atmosphere was re-established. The DIC concentrations in the water were calculated based on temperature, salinity and pH level of the water using the *Seacarb* package in R (Team, 2013). The DIC data was plotted against time.

Based on the outcome of the experiment, the increased average DIC level was calculated for the number of ships passing a 1×1 km grid during 24 hours. In this model, we assumed 1) all CO_2_ produced by a ship is emitted underwater and immediately dissolves; 2) the background and saturated DIC levels of the local water are the same as in the laboratory test, 2.07 mmol/l and 3.97 mmol/l respectively; 3) the temperature condition is the same as in the laboratory test, 20°C, and 4) after a ship passed, the DIC concentration returns to the atmospheric equilibrium in the same time as in the laboratory test. The relationship between the average increased DIC and the number of ships in 24 hours was applied to the ship traffic dataset for creating a global DIC exposure map.

In the actual shipping condition, the CO_2_ will be injected along the ship’s hull instead of in the entire grid cell (1 km x 1 km). Therefore, the actual size of the exposure area was estimated by assuming 1) the injected CO_2_ impacts a 5m depth of water column, and 2) the DIC saturation level (3.97 mmol/l) has to be reached before additional CO_2_ could impacts a larger waterbody. Finally, the impacted surface area (DIC saturated area) of a grid cell was calculated by dividing the DIC saturated water volume by the impacted depth (5 m).

### 2.3 Vulnerability assessments of ecoregions

#### 2.3.1 Vulnerability for CO2 induced algal blooms

From the collected chlorophyll-a concentration dataset, 5000 random data points were plotted against their background DIC levels (Figure 2). The plot shows no relation between DIC and chlorophyll-a at DIC levels above 1840 μmol/kg. At lower DIC levels, the lower limits of the plot show a negative correlation between DIC and chlorophyll-a. This suggests depletion of DIC by primary production, thus a situation where additional CO_2_ can facilitate further algal development. As 1 mg/m^3^ chlorophyll-a is considered as the low limit of a “productive area” for microalgae (Demarcq et al., 2007; Nixon & Thomas, 2001), situations with < 1 mg chlorophyll-a /m3, that only occur at DIC > 1840 μmol/kg are unlikely to be limited by DIC. This condition was therefore assigned as having low vulnerability for CO_2_-induced algal blooms (score 0 - 0.2) (Table 2). In order to derive a relative vulnerability score for situations with Chlorophyll-a > 1 mg/m3, the background DIC levels of the 5000 random data points (Figure 2) were divided into 4 ranges with DIC = 1840 μmol/kg as median value. Vulnerability scores for chlorophyll-a > 1 mg/m^3^ situations were then defined as “0.2 - 0.4”, “0.4 – 0.6”. “0.6 – 0.8” and “0.8 - 1”, when 1060 < DIC < 1460 μmol/kg, 1460 < DIC < 1840 μmol/kg, 1840 < DIC < 2100 μmol/kg and 2100 < DIC < 2347 μmol/kg, respectively (Table 2). Here, a range of vulnerability scores was given instead of a categorical score, because continuous values of vulnerability generate a map with gradual changes, which better represents reality. Within each vulnerability range, a high vulnerability score was given to the place with a relatively low background DIC level as a relatively small addition could be enough to facilitate an algal bloom in a situation with enough nutrients.

**Table 2.** Relative Vulnerability Classes for Elevated CO_2_ Concentrations Assigned to Local Water Conditions Indicating Sensitivity for CO_2_ Induced Algal Blooms or Acidification.

|  | Vulnerability range and conditions |  |  |  |  |
| --- | --- | --- | --- | --- | --- |
|  | 0 - 0.2 | 0.2 - 0.4 | 0.4 - 0.6 | 0.6 - 0.8 | 0.8 - 1 |
| Chlorophyll-a<br>(mg /m<sup>3</sup>)<br>& DIC (µmol/kg) | <1 &<br>1840 -<br>2347 | >1 &<br>2100 -<br>2347 | > 1 &<br>1840 -<br>2100 | > 1 &<br>1460 -<br>1840 | > 1 &<br>1060 - 1460 |
| TAlk (µmol/kg) | 2450 -<br>2400 | 2400 -<br>2350 | 2350 -<br>2300 | 2300 -<br>2250 | 2250 - 2200 |

*Note. The criteria include the chlorophyll-a concentration (mg/m<sup>3</sup>) as a measure for nutrient availability, dissolved inorganic carbon (DIC) concentration (µmol/kg), and Total Alkalinity (TAlk) levels (µmol/kg).*

**Figure 2.**
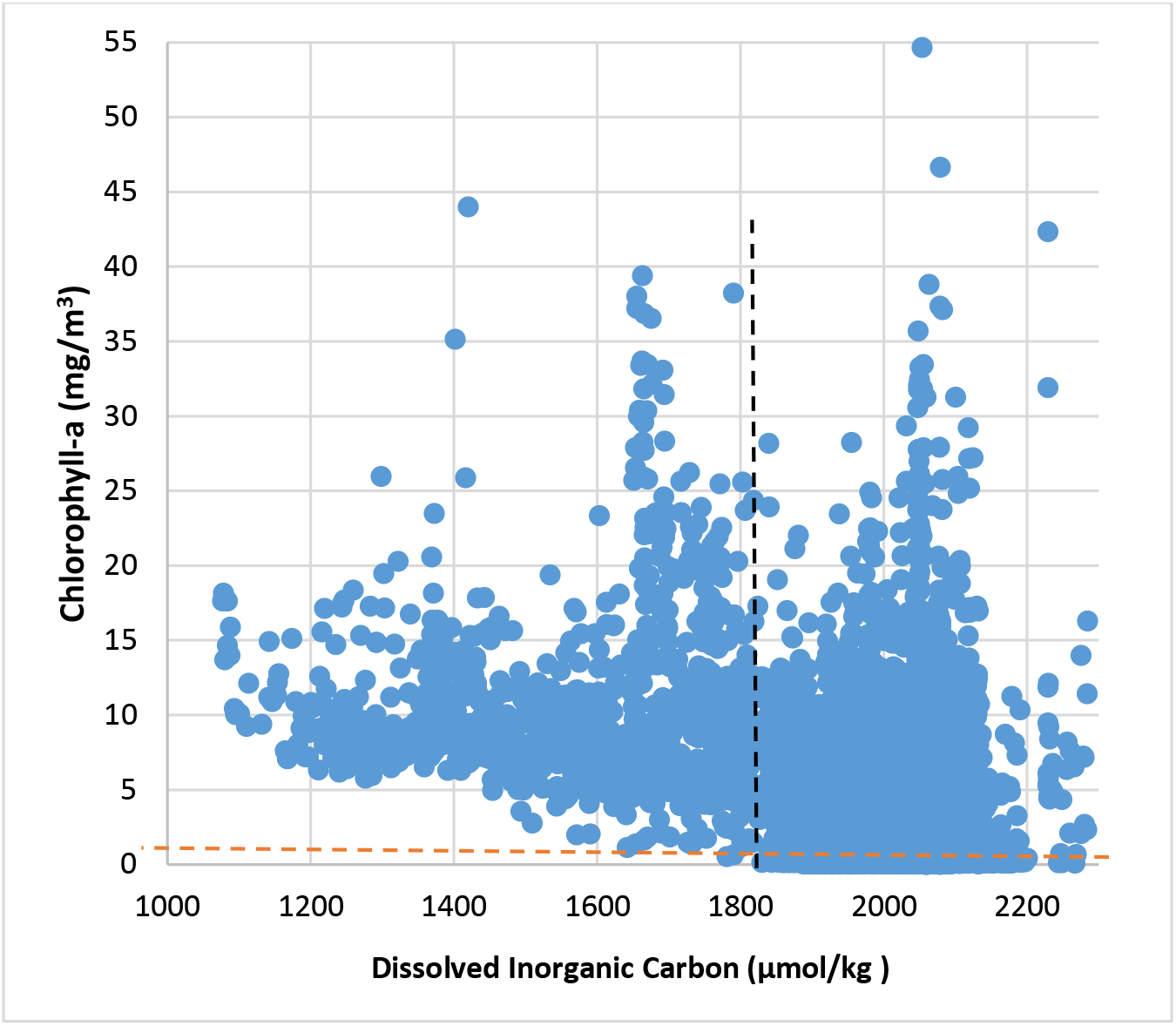
Chlorophyll-a concentration (mg/m^3^) and the yearly average DIC level (μmol/kg) of 5,000 random sample points of the NASA’s Aqua-MODIS satellite dataset in 2018 (NASA Goddard Space Flight Center, 2018). Sample points with 1 mg chlorophyll-a/m^3^ and DIC with 1840 μmol/kg are indicated as red and black dash line, respectively. 1 mg/m^3^ chlorophyll-a was reported as the threshold value to define the low limits of a “productive area” for microalgae (Demarcq et al., 2007; Nixon & Thomas, 2001). The linear correlation between chlorophyll-a concentration and DIC level is not observed when DIC > 1840 μmol/kg in this figure.

#### 2.3.2 Vulnerability to acidification

The derived monthly TAlk values were aggregated to seasonal mean (same season as chlorophyll-a concentration) levels. A TAlk level around the global mean, 2300 - 2350 μmol/kg, was classified as ‘medium’ vulnerability to acidification with a score range “0.4 – 0.6”. TAlk values below this range have a lower acid-neutralizing capacity and therefore the vulnerability score increased with increments of 50 μmol/kg to a maximum score of 1.0 at TAlk 2200 μmol/kg. The same approach was followed in the opposite direction, where the vulnerability score was reduced with increments of 50 μmol/kg to a minimum vulnerability score of 0 at 2450 μmol/kg (Table 2).

#### 2.3.3 Vulnerability assessment

The vulnerability to algal bloom (chlorophyll-a and DIC) and acidification (TAlk level) were given equal weights (0.5). The final per-pixel (1×1 km) vulnerability score (*Overall Vuln*) could be computed by summation of the weight (*0.5*) of each indicator (TAlk and Chlorophyll combined with background DIC level (Chlo. _DIC)) multiplied by the vulnerability score (*S*) of each indicator, as:

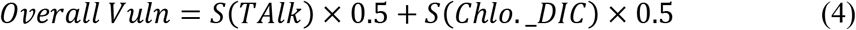

### 2.4 Relative risk assessment

The relative risk level of each ecoregion to acidification and algal bloom was characterized by plotting the DIC exposure level via maritime shipping against the vulnerability score. The ecoregions with relatively high increased DIC concentrations and higher vulnerability scores are considered as higher risk than the ecoregions with lower increased DIC concentrations and vulnerability scores.

The 262 global ecoregions and 144 European sub-ecoregions were mapped and assigned with code numbers. On this map, the layer with the average DIC exposure level via maritime shipping and the average seasonal vulnerability to algal bloom and acidification of each region were mapped. Also, the specific DIC exposure level and vulnerability scores for algal bloom and acidification of individual ecoregion can be found via their code numbers.

## 3. Results

### 3.1 Ship intensity and emitted CO_2_

The worldwide total CO_2_ emission via marine transportation in 2018 was estimated at 1,389 million tonnes under a worst-case-scenario (based on the upper emission factor from Table 1) (Figure 3). The emissions per grid cell were relatively low in the open sea due to the high dispersion of shipping lanes. Contrastingly, high emissions per grid cell were found at dense shipping lanes in coastal areas. Between 1,549 and 12,734 tonnes of CO_2_ were emitted per grid cell in the top 5% busiest ship areas (Figure 3). The highest CO_2_ emission per grid cell was estimated for the Strait of Gibraltar at 12,734 tonnes, closely followed by other maritime chokepoints such as the Panama Canal, the Malacca Strait, the Strait of Hormuz and the Danish Straits (Figure 3).

**Figure 3.**
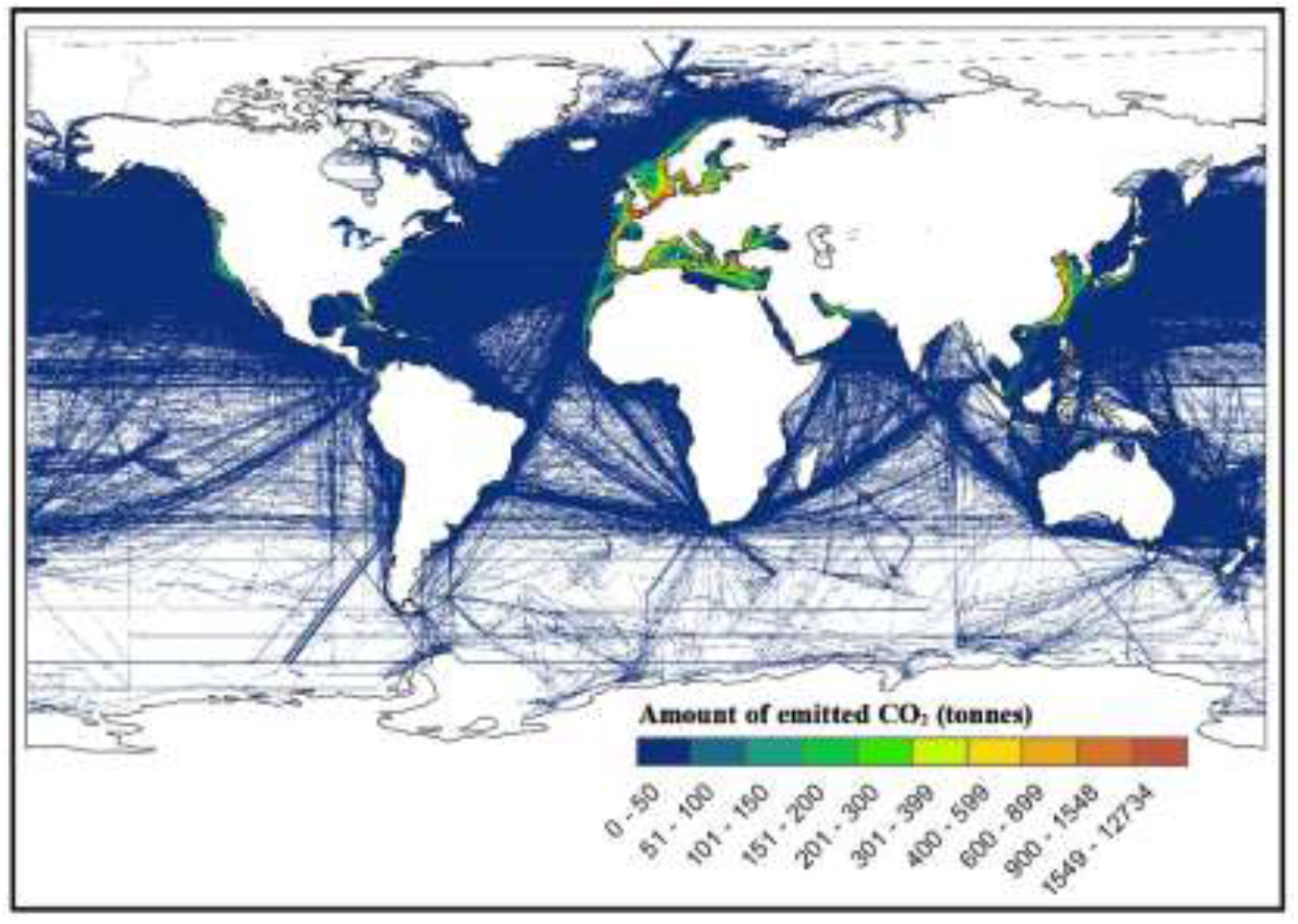
The estimated total amount of CO_2_ (tonnes) emitted from shipping in 2018. The estimation was based on a worst-case-scenario by using the upper emission factors from Table 1. Colours indicate the estimated CO_2_ amount emitted in each 1 km^2^ grid cell.

### 3.2 DIC exposure assessment

The background and saturated DIC concentrations in this study were set at 2.07 mmol/l and 3.97 mmol/l respectively and the temperature was set at 20°C. Thus, a maximum of 1.90 mmol/l DIC can be added via CO_2_ exposure. The estimated added DIC concentration steeply went up with increasing number of ships per 24 hours until after the ninth ship the concentration reached 1.61 mmol/l, so still below the maximum level of 1.90 mmol/l) (Figure 4). From then on, the added DIC concentration levelled off at around 1.61 -1.90 mmol/l regardless of the shipping frequency. Therefore, we assumed that after 9 ships, more volume of water started to experience an increase in the DIC level. With less than 9 ships passing by, the volume of impact water remained ≤ 17 m^2^ of the grid cell (Table 3). In the highest shipping intensity grid cell (134 ships/24 hours), 228 m^2^ water of the 1 km^2^ grid cell (0.023% of the grid cell) reached a saturated DIC level.

**Table 3.** The Relationship Between the Number of Ships (nr. ships) and Size of DIC Saturated Area.

| Ships nr. | $\leq 9$ | 10 | 20 | 30 | 60 | 90 | 120 | 134 |
| --- | --- | --- | --- | --- | --- | --- | --- | --- |
| DIC saturated area (m<sup>2</sup>) | $\leq 17$ | 19 | 34 | 51 | 102 | 153 | 204 | 228 |
| % of 1 km<sup>2</sup> grid cell | 0.002 | 0.002 | 0.003 | 0.005 | 0.010 | 0.015 | 0.020 | 0.023 |

*Note. The table shows the number of ships passing through a 1km<sup>2</sup> grid cell during 24 hours and the part (total m<sup>2</sup> and %) of the 1 km<sup>2</sup> grid cell that will be completely DIC saturated. The total impacted area will be larger (not fully DIC saturated).*

**Figure 4.**
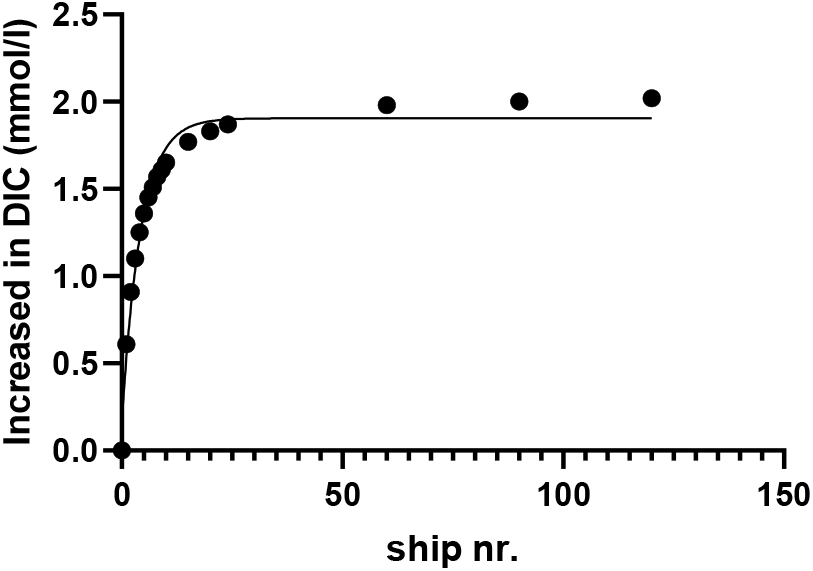
The correlation of increased in DIC level (mmol/l) and the number of ships (ship nr.) passing through a 1 km^2^ grid cell during 24 hours. Assumptions: 1) all CO_2_ produced by the ship(s) is emitted underwater and immediately dissolves and 2) the background and saturation DIC levels of the local water are 2.07 mmol/l and 3.97 mmol/l, respectively, therefore the maximum increase in DIC level is 1.90 mmol/l.

Approximately 70% of the grid cells (with registered ships) in the open sea experienced a DIC increase between 0.15 - 0.22 mmol/l, corresponding to less than one ship per 24 hours, since 1 ship/24 hour already can increase 0.51 mmol/l DIC level (Figure 5). High average DIC exposure (0.52 – 1.9 mmol/l, were located along with European, Chinese and North American coastlines. Eventually, ecoregions along European coastlines were selected to run a zoomed-in local relative risk assessment (section 3.4).

**Figure 5.**
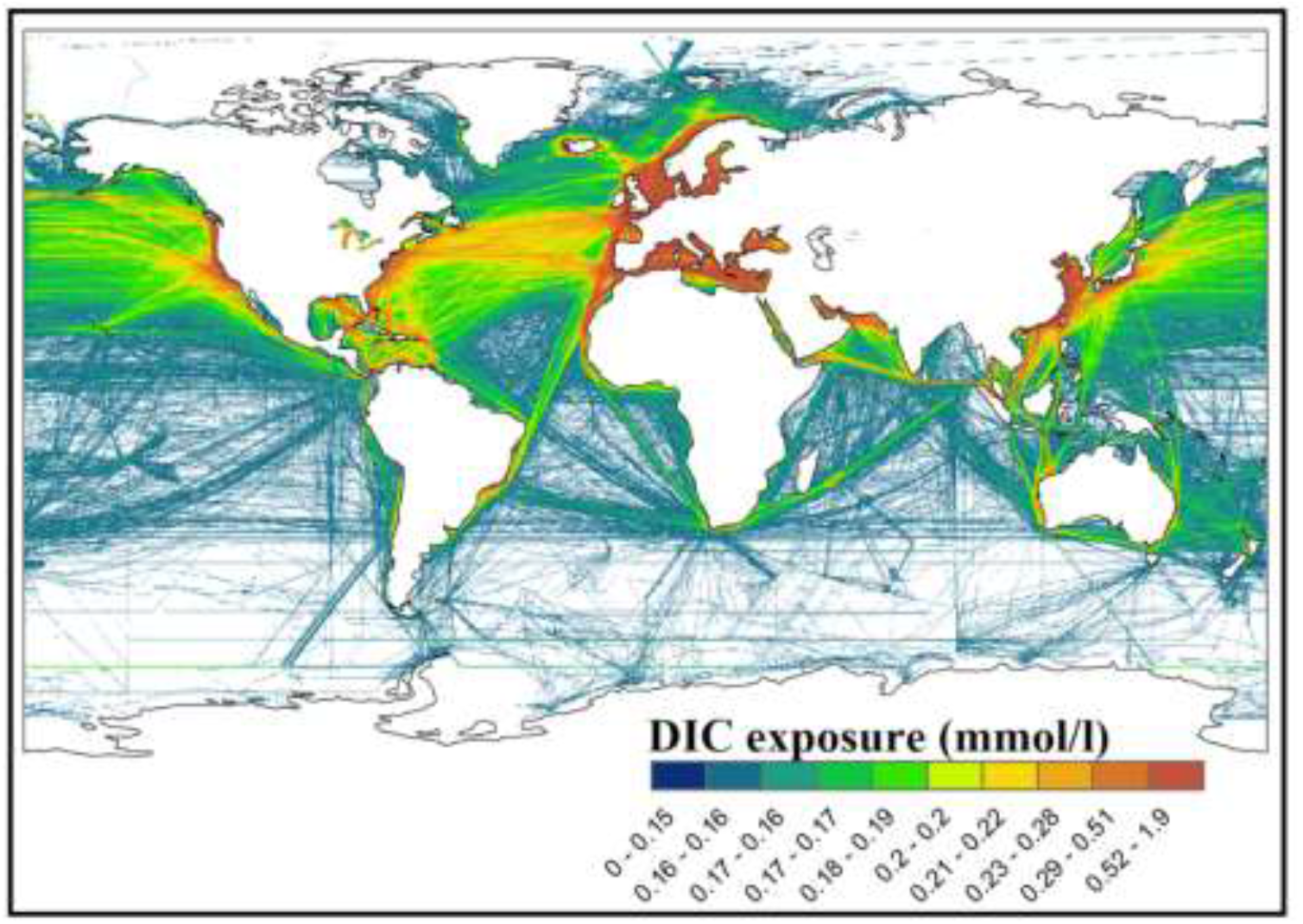
Estimated dissolved inorganic carbon (DIC) exposure (mmol/l) per 24 hours from global shipping, assuming 100% underwater exhaust emission. The scale maximum is restricted by the maximum dissolved CO_2_ saturation level. Colours indicate DIC exposure level in each 1 km^2^ grid cell.

### 3.3 Vulnerability assessment

The spatial distribution of global ocean vulnerability to algal blooms in 2018 was studied based on chlorophyll-a concentration with the background DIC level, and its vulnerability to acidification was evaluated based on the TAlk level. For the “chlorophyll-a and background DIC”, almost all vulnerability score ≥ 0.5 areas were located around the coastal lines of the Northern Hemisphere (Figure S2 and S3 in the supporting information). Contrastingly, almost the entire Southern Hemisphere showed a vulnerability score ≤ 0.2, except the coastline along with Argentine with scores between 0.5 - 0.6 (Figure S2 and S3 in the supporting information). For the TAlk level, areas with > 2500 μmol/kg TAlk were found in the Atlantic Ocean (Figure S4 in the supporting information). While the Pacific Ocean, especially along its coastlines, showed a lower TAlk level (higher vulnerability) compared with other oceans. More seasonal variation seems to occur in the Northern Hemisphere than it does in the South.

After combining all the vulnerability scores, the high vulnerability areas to both acidification and algal blooms were mainly found in coastal areas and above 30° N latitude till the ice edge (Figure 6). The sea ice areas (polar zone) are not being considered in discussion and conclusion due to the missing data in the input datasets (Figure 6). A seasonal impact on the results was observed within the same region, with higher vulnerability scores in the warm season than for the cold season (Figure 6 and Figure S5 in the supporting information).

**Figure 6.**
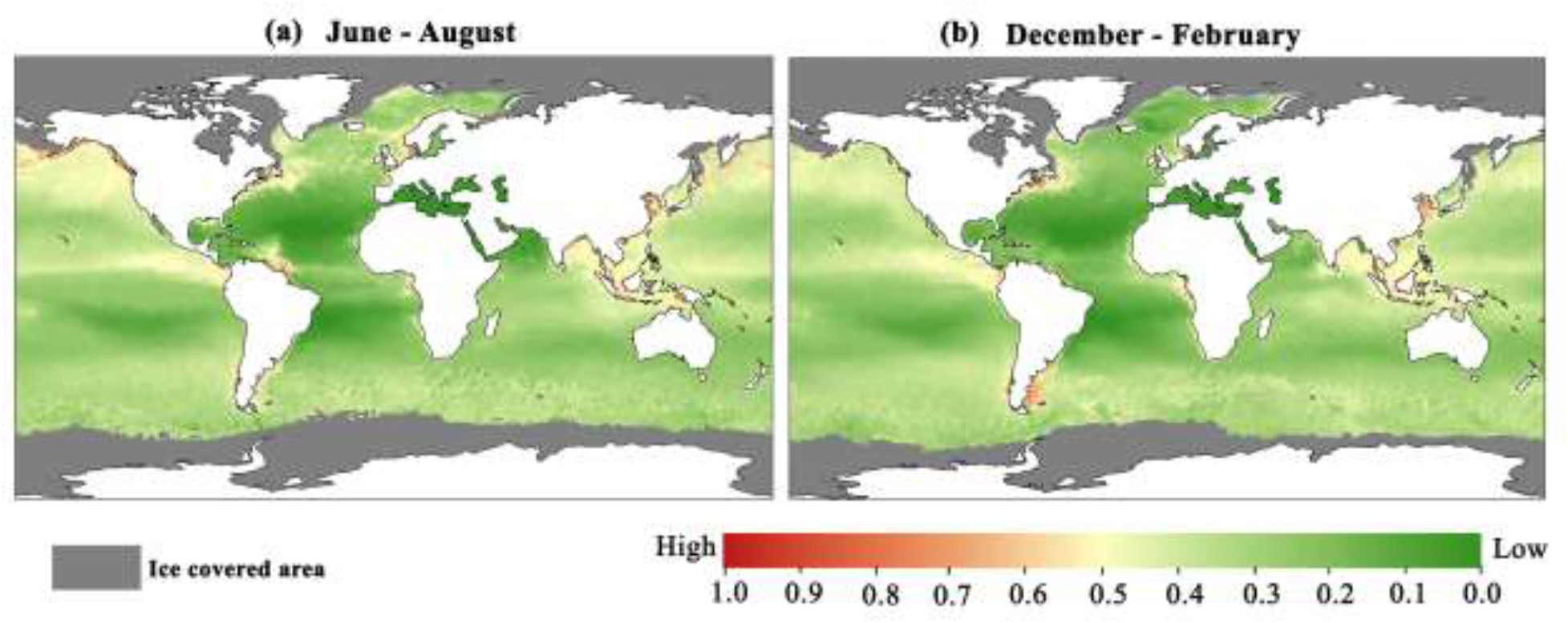
Spatial distribution of vulnerability to algal blooms and acidification in surface layers of the global oceans in June - August (**a**), and December - February (**b**) of 2018. Colours show gridded values based on a merge of three vulnerability indicators, chlorophyll-a & DIC and TAlk as presented in Table 2.

### 3.4 Relative risk assessment

Over 90% of the global ecoregions showed a DIC exposure level < 0.48 mmol/l and a vulnerability score to acidification and algal blooms < 0.5 (Figure 7). The Yellow Sea and Southern China Sea showed relatively higher risk than other areas in all seasons (Figure 7, Figure S6, S7 and Table S1 in the supporting information). Next in line is the North Sea, which is exposed to over 0.95 mmol/l DIC by maritime emissions, and a relatively high vulnerable score close to 0.5 especially during June – August. Seasonal variation in shipping intensity was not included in this study. Therefore, the DIC exposure level of each region was consistent over the year. The seasonal variation of global risk was therefore only influenced by the seasonal vulnerability score (Figure 6 and Figure 7).

**Figure 7.**
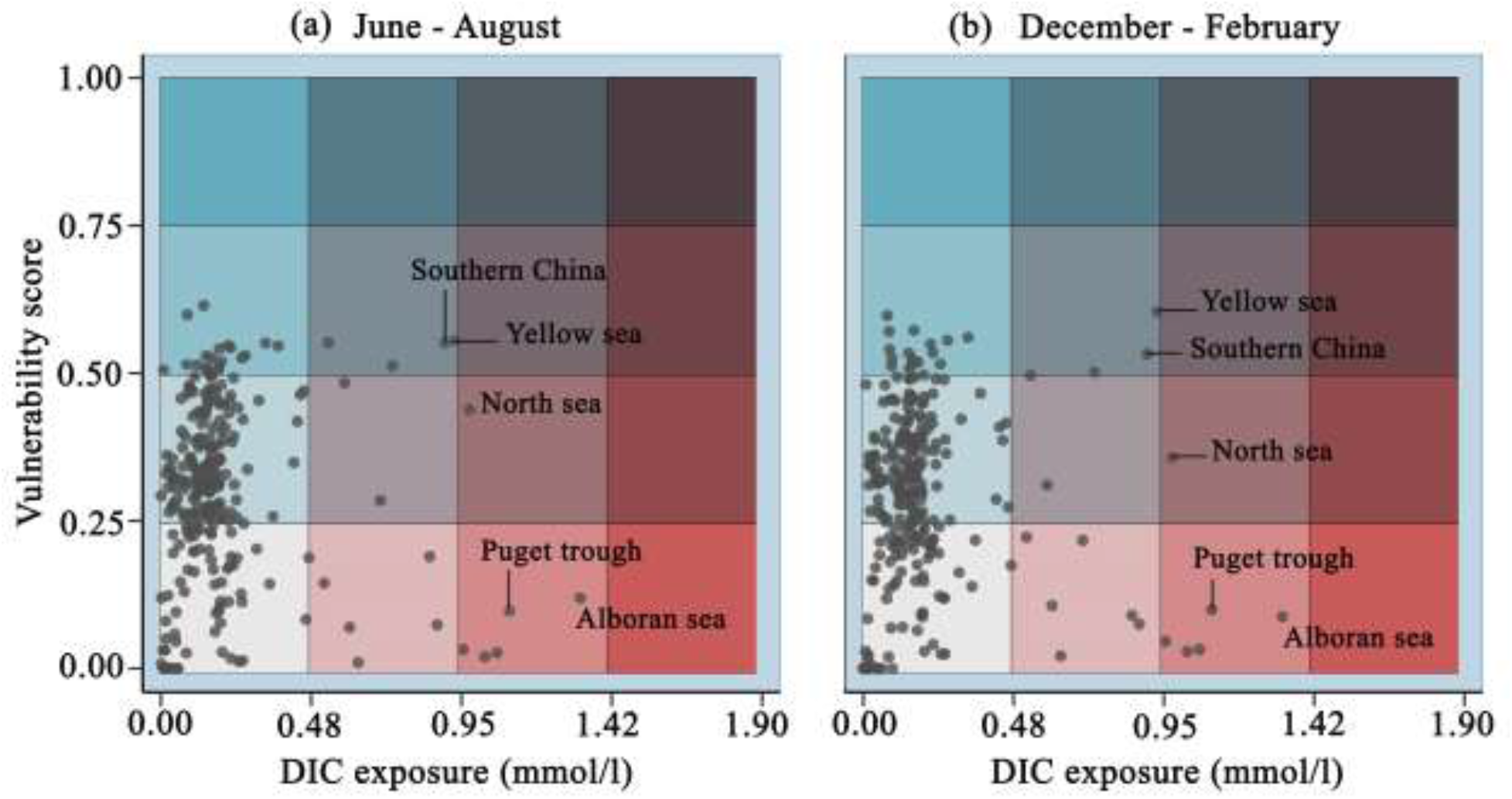
Plot of the vulnerability scores of 262 global maritime ecoregions to algal blooms and acidification and the estimated DIC exposure (mmol/l) by shipping in June – August (**a**) and December – February (**b**). Assumption: all ships would be equipped with underwater exhaust systems. The colour intensity indicates the vulnerability score (blue) and increase in DIC level (red) combined into a relative risk level from low (light grey) to high (blue/red).

When looking at a more detailed level to Europe, it becomes clear that the relatively high risk that was predicted for this ecoregion only concerns areas with dense shipping lanes and maritime chokepoints, such as the Strait of Dover and the Strait of Gibraltar (Figure 8). The biggest seasonal increase in the relative risk could be observed in coastal areas of the Celtic Seas, the Saharan Upwelling, and the South European Atlantic Shelf in spring and summer (Figure 8, Figure S8, S9 and Table S2 in the supporting information).

**Figure 8.**
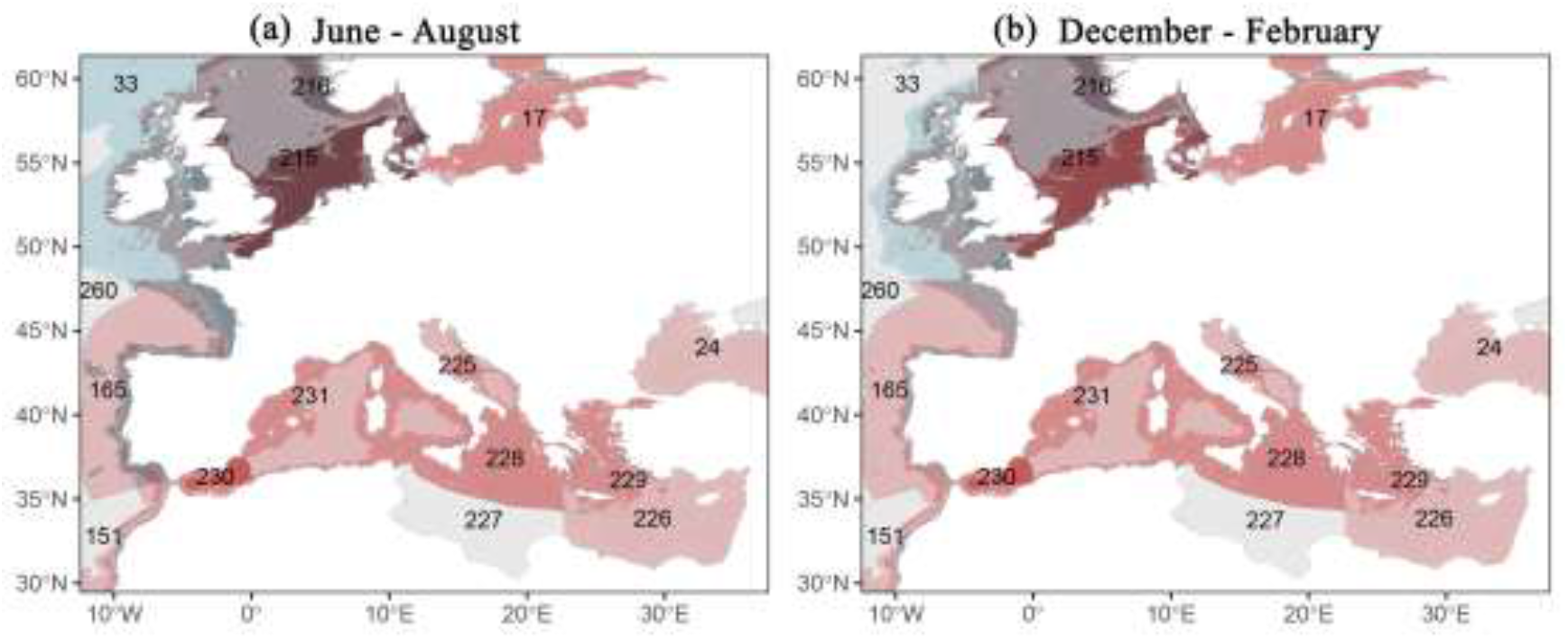
Spatial distribution of the relative risk for algal blooms and acidification in the 15 European marine ecoregions (global ecological codes: 17, 24, 33, 151, 165, 215, 216, 225, 226, 227, 228, 229, 230, 231, and 260 (Mark D. Spalding et al., 2007b) in June – August (**a**), and December – February (**b**). Colours show gridded values from plotting vulnerability scores against DIC exposure caused by maritime shipping emitting CO_2_ underwater. The vulnerability score and DIC exposure level of each ecoregion are presented in the separate table (Table S2 in supporting information). The colours in this map are corresponding with the colours in Figure 7. Thus, the colour intensity indicates the vulnerability score (blue) and increase in DIC level (red) combined into a relative risk level from low (light grey) to high (blue/red).

## 4 Discussion

In the presented study an assessment was made of the relative risk that a local marine environment is negatively impacted by the application of underwater released exhaust gas as ‘air lubrication’ along the ship’s hull. High nutrient availability potentially inducing algal blooms or low buffering capacity potentially resulting in acidification were used as indicators of environmental vulnerability. Following a worst-case exposure scenario, it is assumed that all ships will be equipped with underwater exhaust systems and that all emitted CO_2_ is absorbed by the water. Based on this a first-tier risk environmental assessment was performed for 262 ecoregions.

### 4.1 Shipping emitted CO_2_ and DIC exposure

The worldwide CO_2_ emission from marine shipping was extrapolated from data of 2013 to be 1,389 million tonnes in 2018. This number can be considered as the worst case, as current developments suggest a lower growth of CO_2_ emission. IMO predicted the CO_2_ emission to grow 50% - 250% (to 1,194 – 2,786 million tonnes) by 2050, but estimated that emission < 900 million tonnes CO_2_ in 2018 (IMO, 2015). In our study, we also used the upper emission factors from Table 1 which also contribute to a worst-case-scenario of the emitted CO_2_ amount. If the amount of shipping emitted CO_2_ would continue to increase in the future as predicted, the DIC exposure (concentration and surface size) via underwater exhaust gas will increase as well. For the amount of emitted CO_2_ in an individual grid cell, our calculation used the average travel distance in that cell (0.7 km). This was not according to a worst-case-scenario, since the longest travel distance to pass a 1 x 1 km grid cell is over its full diagonal, so 1.41 km. Using the average travel distance in the calculation leads to the amount of CO_2_ emission per grid cell closer to the real conditions, especially from the ecoregion perspective, which consists of many grid cells.

In this study, we excluded the effect of water currents, and thus the transport and dilution of the dissolved CO_2_ that result from that. It can therefore be assumed that our calculations overestimate the volumes of water where the maximum area DIC saturation level will be reached while we underestimate the volume of water that is exposed to lower (diluted) CO_2_ concentrations. We also assumed that all emitted CO_2_ will be completely dissolved in the water, whereas in reality it may be expected that some will escape to the atmosphere. The maximum DIC exposure levels calculated here are worst-case estimations. These and several other aspects need to be taken into account in further refinement of the exposure assessment for areas where the DIC levels are indicated to become a potential problem. Especially the size of the impacted area with water currents, the ship’s speed, the water mixing depth (here set at 5 m depth) and the extent to which the underwater released exhaust CO_2_ is likely to fully dissolve in the water or will partially immediately escape to the atmosphere with gas bubbles. The dissolving of CO_2_ gas in water phases involves a series of reactions (Zeebe & Wolf-Gladrow, 2001), which takes time and has a saturation level assuming a static situation.

For refining the exposure assessment, also the assumed DIC saturation level should be further fine-tuned. The current exposure assessment is based on a single water condition (20 °C, 31.7‰ salinity and 2.07 mmol/l alkalinity) without primary production. The background and saturation level of DIC, however, are influenced by the water conditions, such as temperature, salinity, and algal productivity (Markou et al., 2014; Zeebe & Wolf-Gladrow, 2001). If a specific water condition is known, a prediction of the DIC exposure level can be made based on the presented approach. For example, between July - August, Chukchi Sea shelves experience relatively high water temperature (−1.5 to + 7 °C) and phytoplankton production. Thus, it is expected that the background DIC level in this area is lower than in the cold season and also than the background DIC level assumed in this study. Indeed, the reported background DIC level in part of this area is even below 0.6 mmol/l DIC in July – August (Bates et al., 2005).

### 4.2 Vulnerability assessment

Areas vulnerable to acidification and algal blooms are mostly the subtropical ecotypes, especially near coastal regions and in warm seasons (e.g. East China seas in June - August and south coastline of Argentina in December – February). This distribution can mostly be attributed to the high vulnerability for algal blooms around nutrient rich nearshore areas (Figure S2 in the supporting information). Algal blooms are shown to exacerbate in eutrophic areas during seasonal warming (Lee et al., 2006; Moore & Abbott, 2000). Anthropogenic nutrients input along the coastline and at large river mouths, e.g. from aquaculture, runoff, sewage and other pointsource pollution, is the main driver to create those eutrophic conditions (Halpern et al., 2015).

Other areas with high vulnerability scores (> 0.7) are mainly located above 30°N latitude. L Q Jiang et al. (2015) reported similar results when they identified the vulnerability of the global oceans to acidification via aragonite saturation state, which decreased toward higher latitude after 40°. They attribute this latitudinal gradient to the water temperature influenced change in TAlk/DIC ratio. In our study, the overall high vulnerability scores above 40° latitudes are mainly caused by the low TAlk level in those regions, thus high vulnerability to acidification as well. TAlk is mainly reflected by SSS (sea surface salinity) changes instead of SST (sea surface temperature) (Lee et al., 2006). Low TAlk usually results from low large influxes of freshwater through ice melting (sea ice edge) and through river outflows (e.g. the Amazon, the Congo River and the Bay of Bengal), or where precipitation exceeds evaporation (Buis et al., 2011). Therefore, TAlk levels generally are lower at higher altitudes and usually show high seasonal variation. Another seasonal variable in TAlk was found in the Northern Hemisphere more than in the Southern Oceans (Fine et al., 2015), namely due to high salinity variability and active water currents that resulted in upwelling of water enriched in alkalinity during winter and autumn (Z P Jiang et al., 2014). Such a difference in seasonal impact between north and south was not found in the overall vulnerability distribution map that combines the TAlk with “chlorophyll -a and background DIC” indicators.

It is important to be aware that the vulnerability of areas may change with time due to changes in anthropogenic activities and also due to climate change, especially towards the current ice-covered areas. Halpern et al. (2015) reported that a significant amount of ice was lost over the 5 years period of their study of the human impact on the world’s oceans, demonstrating that the water conditions near the polar zone are rapidly changing with time. Likewise, the estimation of vulnerable areas in the present study will be influenced by such large scale changes.

### 4.3 Potential impact assessment of underwater released exhaust CO2

Ecoregions with a high estimated DIC exposure and vulnerability to algal blooms and acidification would be at risk according to our tier 1 relative risk assessment when all ships would be equipped with underwater exhaust systems. Globally, the Yellow Sea and the Southern China Sea were identified as the ecoregions with the relatively highest risk (relatively high exposure and vulnerability) in those 262 ecoregions, closely followed by the North Sea. There clearly would be hotspots of exposure in the busiest shipping traffic grid cell (134 ships/24 hours) in the Yellow Sea, Southern China Sea and the North Sea.). Based on this first tier potential impact assessment it cannot yet be concluded whether there will be a relatively small area with very high exposure or a larger area with a lower exposure but a greater total DIC increase. But in general terms, it is clear that in these ecoregions the relative risk via high exposure concentration and vulnerability score is high.

The European marine ecoregion risk assessment revealed high local exposure conditions. The result supports the conclusion of the global relative risk assessment that especially dense shipping lanes and maritime chokepoints determine the potential impact of the entire ecoregion. In the more detailed relative risk assessment, seasonal variation in risk was more apparent than in the global assessment. The seasonal variation in the European ecoregions can be attributed to the increased algal density during the warming period (March-August) (Lee et al., 2006; Moore & Abbott, 2000), as well as elevated TAlk concentrations in the North Sea and the Celtic Seas from February through May. The result suggests that risk assessment on smaller ecoregions can reveal specific local conditions that would be unnoticed at the global ecoregions level.

Further fine-tuning still can be achieved for the vulnerability assessment and potential impact assessment. For example, both global and local risk assessments in this study assume a linear response of the ecosystem to increased exposure level and vulnerability scores. However, most marine ecosystems exhibit synergistic and antagonistic responses to stressors instead of additive (Crain et al., 2008), which creates a nonlinear relationship of risk to exposure and vulnerability level. Also, the biological composition of each ecoregion would be relevant to consider, something we did not include in this study. Some nearshore locations dominated by coral reefs, such as the Australian Great Barrier reef, scored low on vulnerability or risk in this study but are quite vulnerable for elevated CO_2_ concentrations (Ainsworth et al., 2016). Therefore, it is recommended to perform local risk assessments for specific ecoregions, also including the unique biological composition and water conditions of the studied regions to identify specific exposure-response relationships.

## 5 Conclusions

In this study, we carried out a first-tier relative risk assessment of potential future underwater released exhaust CO_2_ from merchant ships on marine ecosystems. The relative risk of 262 marine ecoregions for enhanced algal blooms and acidification based on specific water conditions was combined with the predicted additional DIC exposure level for each ecoregion from the extra CO_2_ exposure. Globally, relatively high-risk ecoregions were mainly located in the Northern Hemisphere, especially along coastlines, such as in the North Sea and Southern China sea. Those regions combine high shipping frequencies with high vulnerability to CO_2_ induced algal blooms and acidification. In this study, worst-case exposure-scenarios were applied, that need to be refined to better assess the impacted water volume and area and the maximum DIC level that could be reached. Furthermore, this study paves the path for ongoing risk assessment of underwater released exhaust CO_2_ when more information becomes available. In addition, this approach could be used for sensitivity assessment of ecoregions for future elevated CO_2_ levels, and can be further refined by including additional important parameters, such as the biological composition of the ecosystem and the magnitude and influence of water mixing. For sure it already indicates areas that deserve further attention.

## Acknowledgments

This research was funded from the project ‘GasDrive: Minimizing emissions and energy losses at sea with LNG combined prime movers, underwater exhausts and nano hull materials’ (project 14504) of the Netherlands Organization for Scientific Research, domain Applied and Engineering Sciences (TTW). The authors wish to thank Mr. Achmad Sahri (MSc) for technical support in using the ArcGIS program.

## Data Availability Statement

The datasets of 232 Marine Ecoregions of the World were collected from The Nature ConservancY′s Geospatial Conservation Atlas via link: https://geospatial.tnc.org/datasets/ed2be4cf8b7a451f84fd093c2e7660e3_0?geometry=58.359%2C-89.110%2C-58.359%2C87.258. The 30 Marine Ecoregions and Pelagic Provinces of the World were collected from UNEP’s Ocean Data Viewer: https://data.unep-wcmc.org/datasets/38. Chlorophyll-a data were collected from NASA’s Aqua-MODIS satellite for each calendar month of 2018 via link:https://oceancolor.gsfc.nasa.gov/data/10.5067/AQUA/MODIS/L3M/CHL/2018/. Sea Surface Salinity (SSS) data were collected from NASA’s JPL SMAP satellite (https://doi.org/10.5067/SMP42-3TMCS). Sea Surface Temperature (SST) data were collected from NOAA’s Coral Reef Watch satellites for each calendar month of 2018 (https://data.nodc.noaa.gov/cgi-bin/iso?id=gov.noaa.nodc:AVHRR_Pathfinder-NODC-L3C-v5.2;view=html). A ship traffic intensity map developed by Halpern et al. (2015) was collected from the Knowledge Network for Biocomplexity (https://knb.ecoinformatics.org/view/doi:10.5063/F19Z92TW).

