## Supplemental data for "Relative Risk Assessment of Ecological Areas with the Highest Potential Impact of Underwater Released Exhaust CO_2_ from Innovative Ships"

### Contents of this file

Text S1
Figures S1 – S9
Tables S1 & S2

### Introduction

Here, we explain the average travel distance of a ship through one grid cell (Text S1) and provide an additional figure to show the mean depth classes of the 15 marine ecoregions of Europe (Figure S1). The figures in the article showed the vulnerabilities of ecoregions to acidification and algal blooms and risk characterization of underwater released exhaust $CO_2$ in June – August and December – February. Here, Figures S2, S4 and S5 show the global chlorophyll-a concentration, total alkalinity and vulnerability of ecoregions to acidification and algal blooms in four seasons, respectively. Figure S3 shows the global yearly average dissolved inorganic carbon (DIC) background concentration. All the information in these additional figures contribute to the plotted relative risk characterization in four seasons (Figure S6 – S9). For each ecoregion, the specific level of DIC, chlorophyll-a concentration and total alkalinity can be found in Table S1 (global ecoregions) and S2 (zooming in to European ecoregions).

**Text S1.** The average travel distance of a ship through a 1 $km^2$ grid cell

Given the large distances that ships travel and the assumption that mariners prefer great circle distances (Halpern et al., 2015), it was assumed that ship routes were straight from their entry point in a grid cell to their exit point in that grid cell. With straight routes, the maximum distance a ship could travel to pass through a grid cell (1 km x 1 km) is the maximum diagonal distance, so about 1.4 km. A random float number between 0 and 1.4 was generated for each vessel (94169 total). The mean of the randomly generated numbers is 0.7, which represented the average distance a ship travels to pass through a grid cell.

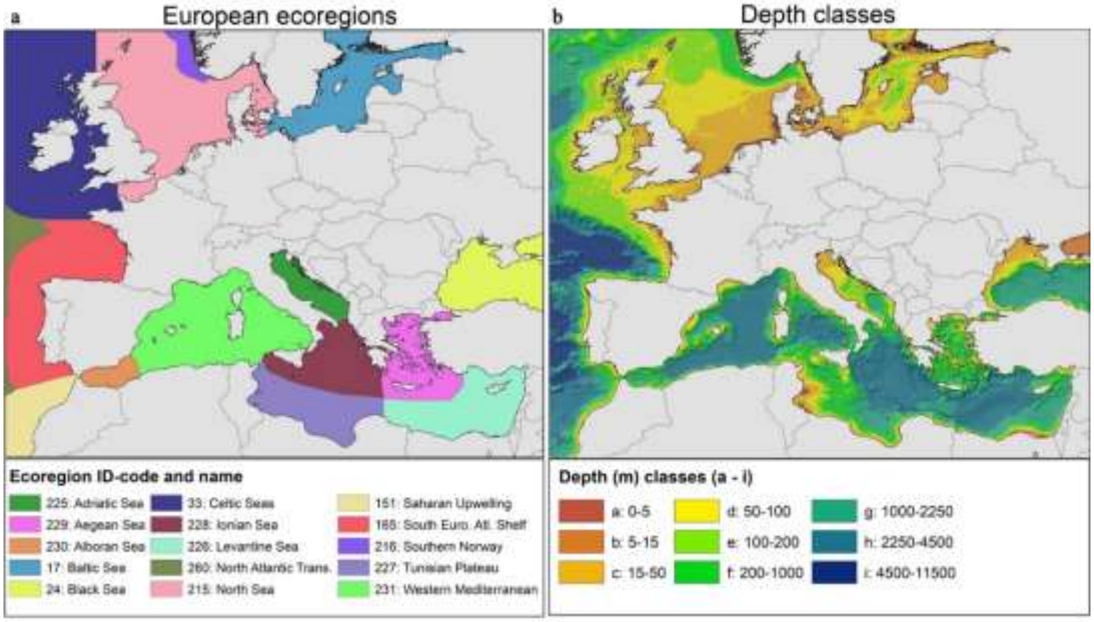


**Figure S1.** The 15 marine ecoregions of Europe (a) and their bathymetry depth classes (b).

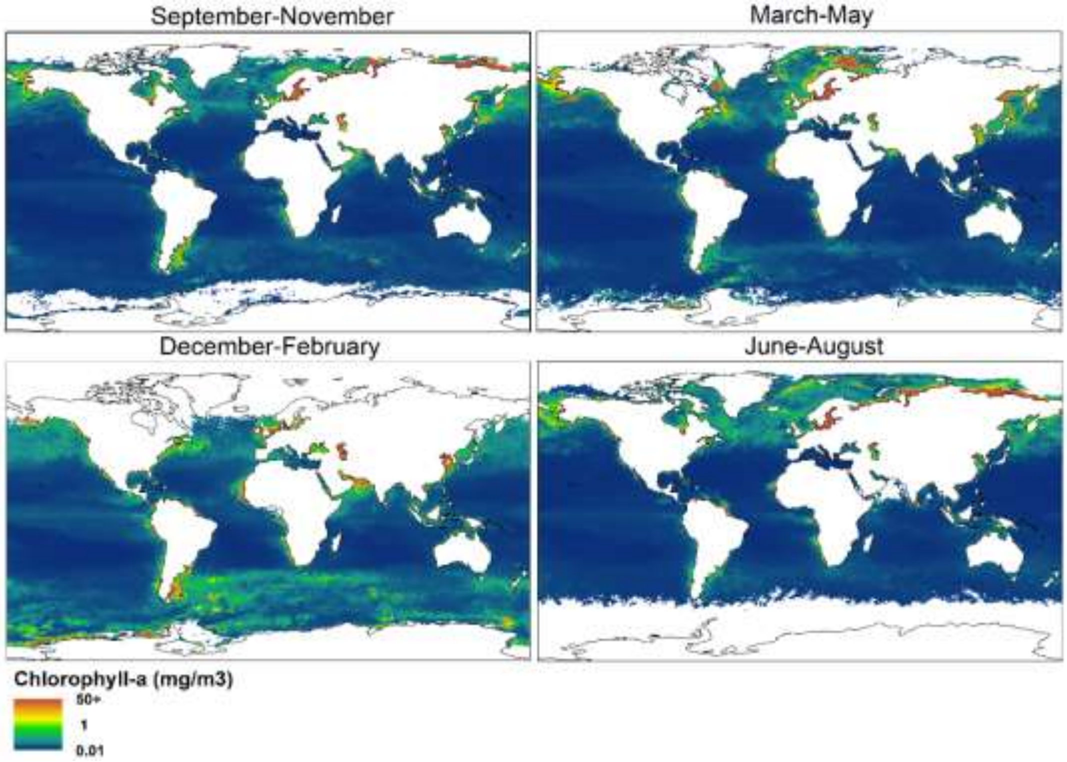


**Figure S2.** Distribution of global chlorophyll-a concentration (mg/m$^3$) in four seasons. Colours indicate the values of the chlorophyll-a concentration in each 1 km$^2$ grid cell.

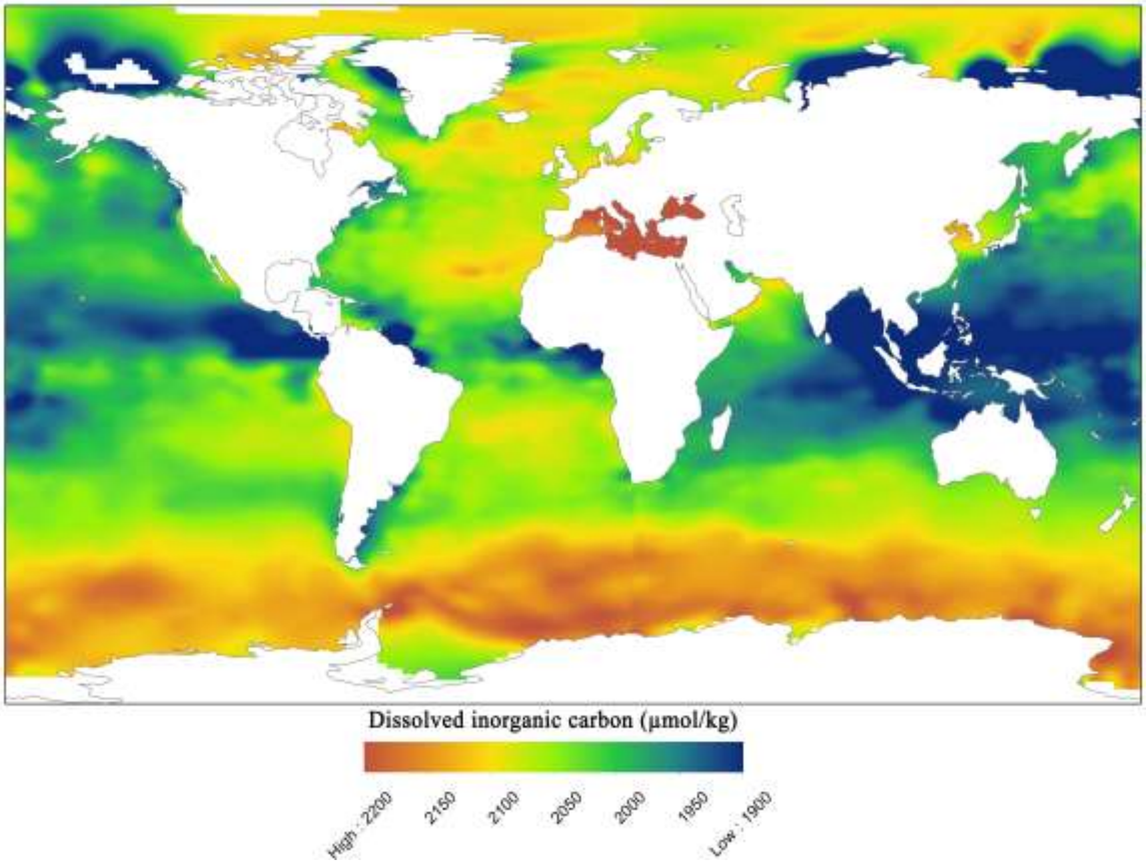


**Figure S3**. The global yearly average dissolved inorganic carbon (DIC) background concentration (µmol/kg) indicated with colours per 1 km$^2$ grid cell.

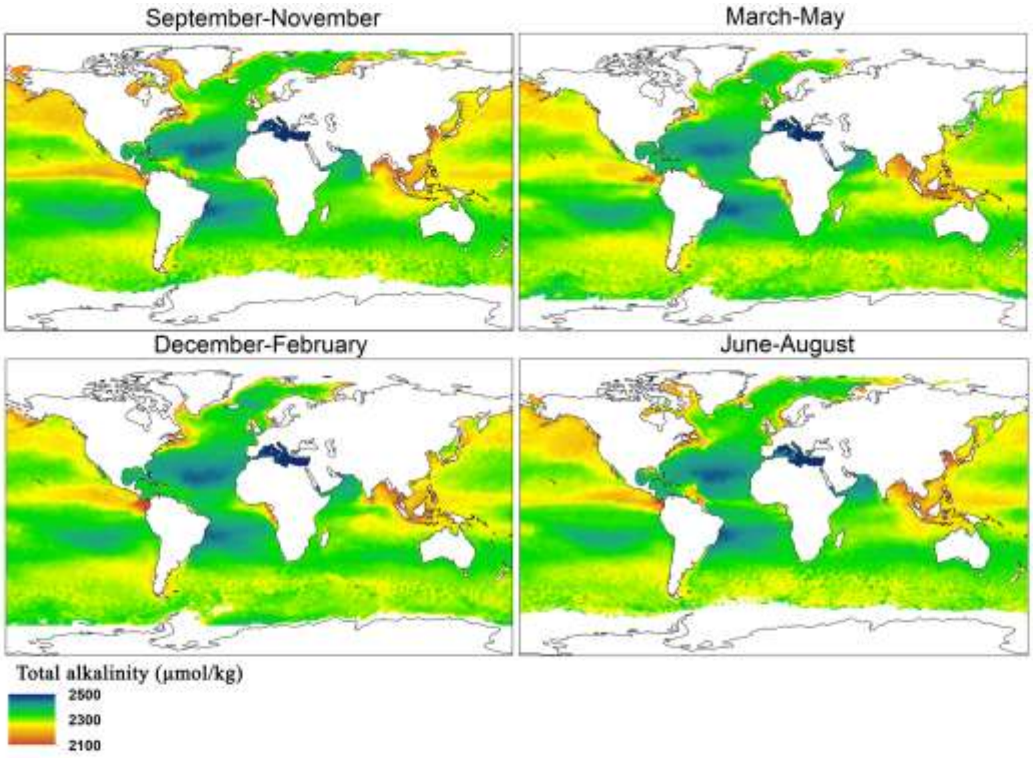


**Figure S4.** Distribution of the global total alkalinity (TAlk) level (μmol/kg) in four seasons, indicated with colours per 1 km$^2$ grid cell.

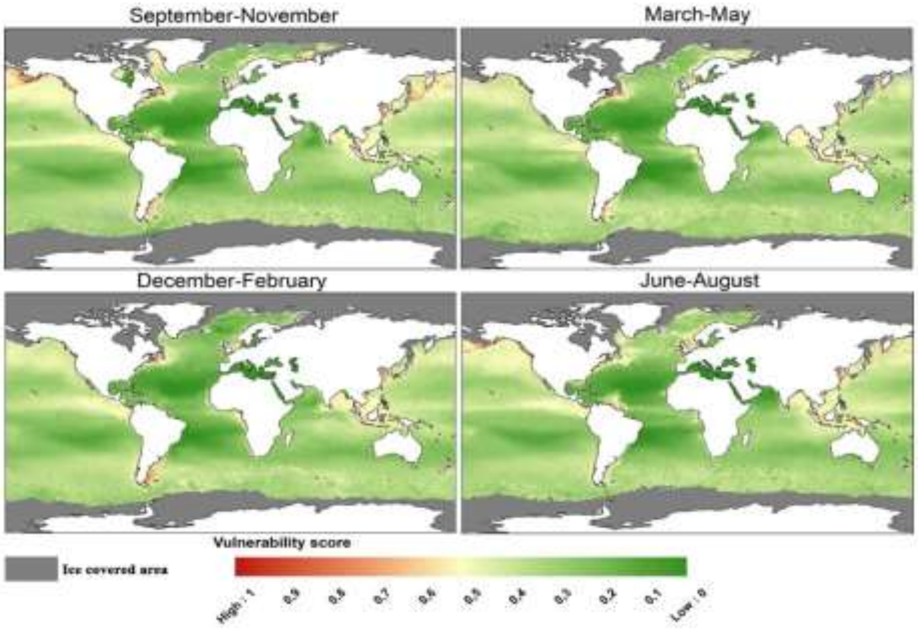


**Figure S5.** Global distribution of vulnerability to algal blooms and acidification in four seasons. Colours show gridded values based on a merge of three vulnerability indicators, chlorophyll-a & background DIC and TAlk.

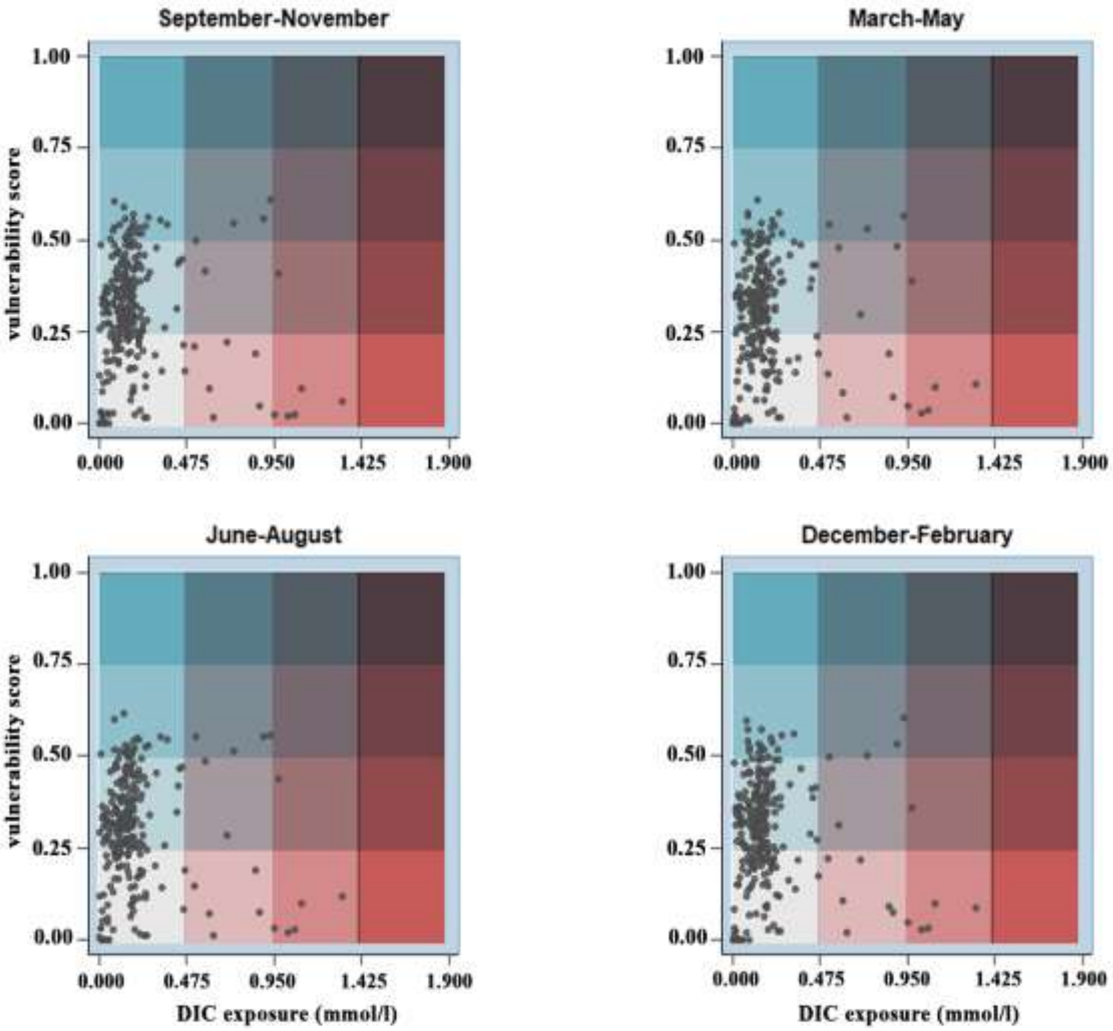


**Figure S6.** Plot of the vulnerability of 262 global ecoregions for algal blooms and acidification following predicted extra DIC exposure (mmol/l) from maritime shipping with underwater released CO2, in four seasons. The colour intensity indicates the vulnerability score (blue) and increase in DIC level (red) and risk from low (light grey) to high (blue/red). The data underlying these plots can be found in Table S1.

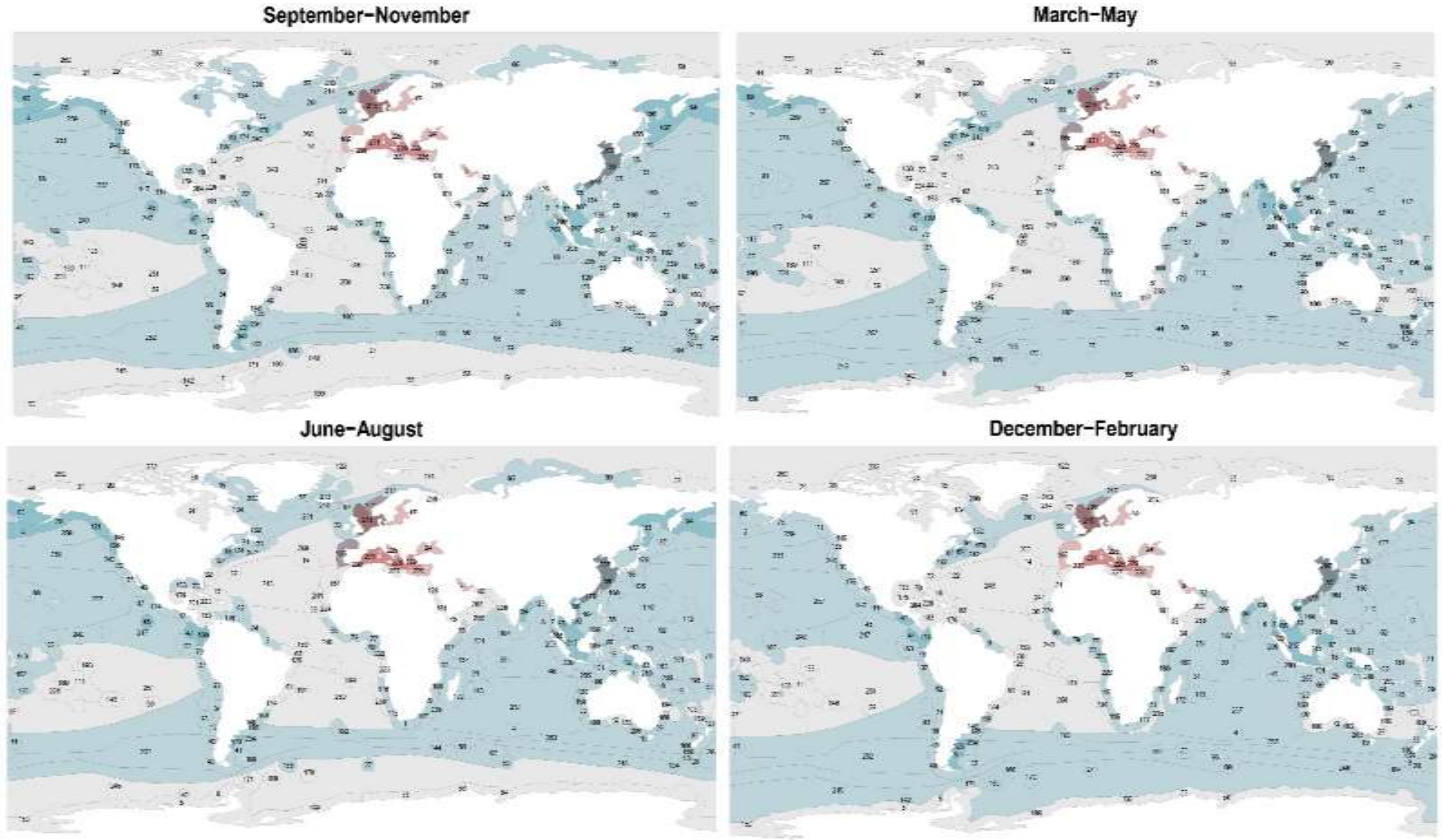


**Figure S7.** Map of the vulnerability of 262 global ecoregions for algal blooms and acidification following predicted extra DIC exposure (mmol/l) from maritime shipping with underwater released $CO_2$, in four seasons.

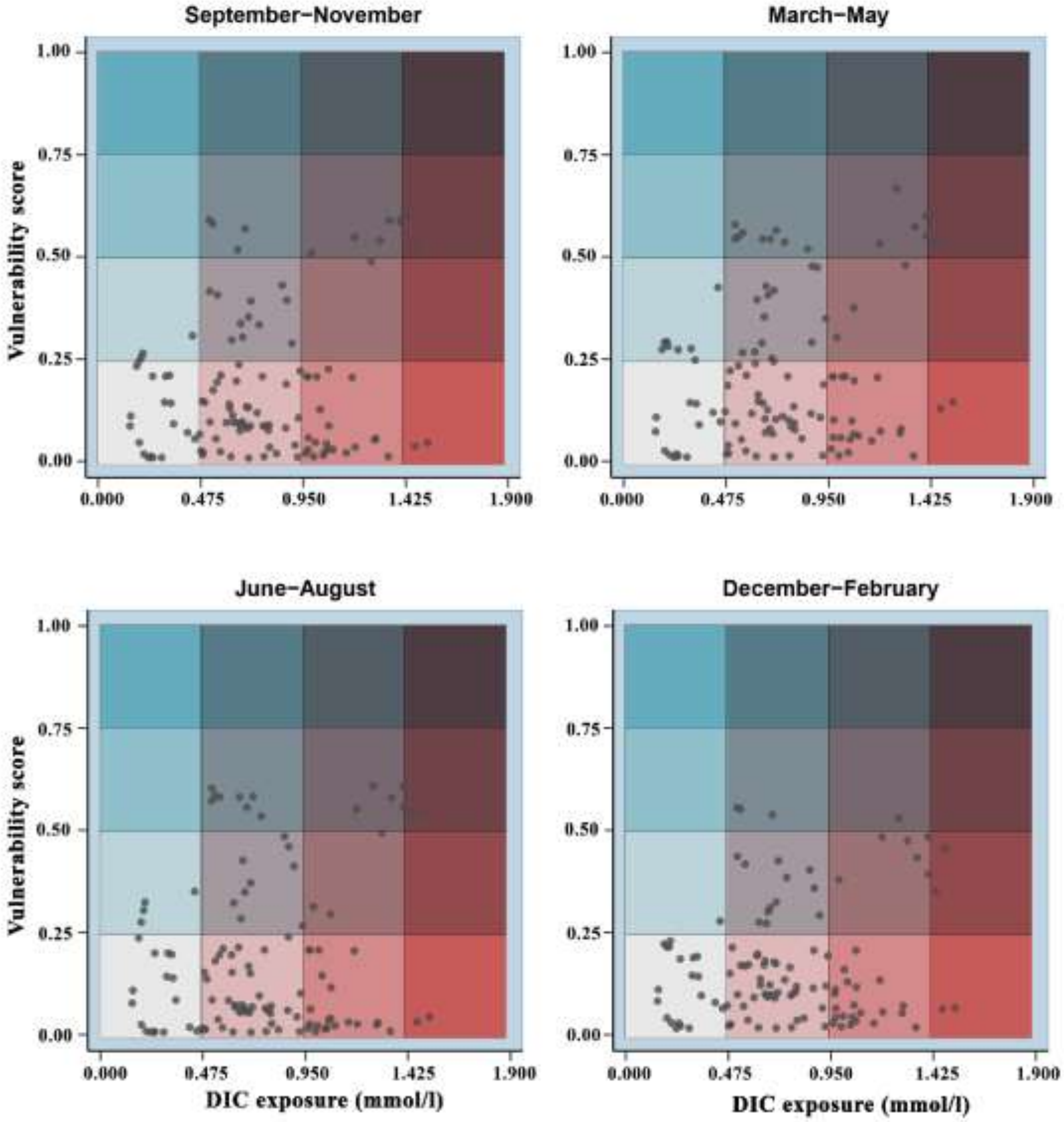


**Figure S8.** Plot of the vulnerability of 115 European sub-ecoregions for algal blooms and acidification following predicted extra DIC exposure (mmol/l) from maritime shipping with underwater released $CO_2$, in four seasons. The colour intensity indicates the vulnerability score (blue) and increase in DIC level (red) and risk from low (light grey) to high (blue/red). The data underlying these plots can be found in Table S2.

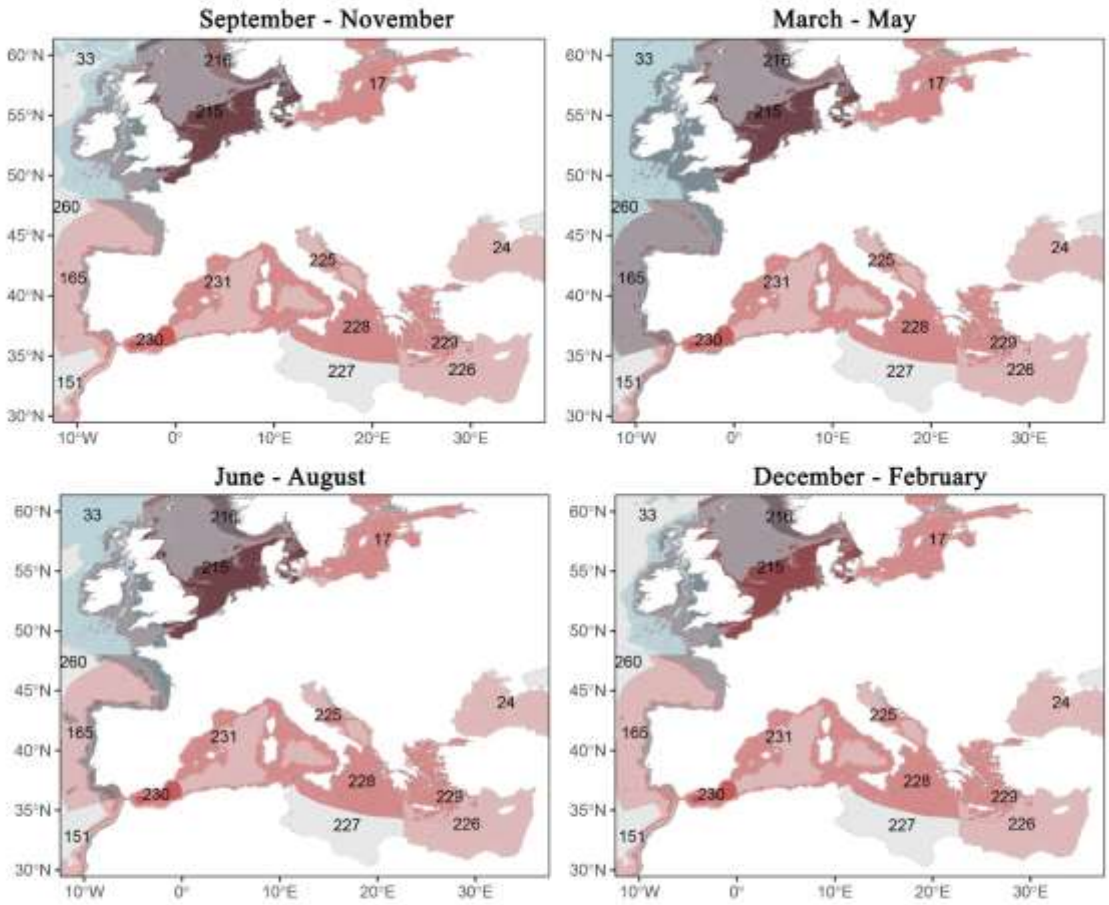


**Figure S9.** Map of the risk of European sub-ecoregions for algal blooms and acidification following predicted extra DIC exposure (mmol/l) from maritime shipping with underwater released $CO_2$, in four seasons.

**Table S1.** Predicted extra DIC Exposure level and vulnerability scores of the 262 global ecoregions in four seasons. The risk for adverse effects from maritime shipping with underwater released CO2, as also presented in Figures S6 and S7, can be calculated by multiplying exposure with vulnerability.

| Code | Ecoregion | DIC exposure (mmol/l) | Vulnerability to algal blooms and acidification | | | |
|---|---|---|---|---|---|---|
| | | | Sep. - Nov. | Mar. - May | Jun. - Aug. | Dec. - Feb. |
| 1 | Agulhas Bank | 0.1821 | 0.3371 | 0.2977 | 0.3153 | 0.3023 |
| 2 | Aleutian Islands | 0.1822 | 0.5338 | 0.4626 | 0.5124 | 0.4496 |
| 3 | Amazonia | 0.1607 | 0.2385 | 0.2768 | 0.2710 | 0.2574 |
| 4 | Amsterdam-St Paul | 0.0479 | 0.2730 | 0.2635 | 0.2713 | 0.2721 |
| 5 | Amundsen/Bellingshausen Sea | 0.0135 | 0.0018 | 0.0407 | 0.0000 | 0.0845 |
| 6 | Andaman and Nicobar Islands | 0.1063 | 0.4714 | 0.5086 | 0.4944 | 0.4935 |
| 7 | Andaman Sea Coral Coast | 0.0904 | 0.3930 | 0.5202 | 0.4756 | 0.5254 |
| 8 | Antarctic Peninsula | 0.0509 | 0.0249 | 0.1829 | 0.0003 | 0.1921 |
| 9 | Arabian (Persian) Gulf | 0.5160 | 0.2117 | 0.1340 | 0.1447 | 0.2223 |
| 10 | Araucanian | 0.1559 | 0.4964 | 0.4960 | 0.4959 | 0.4587 |
| 11 | Arnhem Coast to Gulf of Carpenteria | 0.1249 | 0.3701 | 0.4866 | 0.4317 | 0.4188 |
| 12 | Arafura Sea | 0.0834 | 0.5334 | 0.5745 | 0.5992 | 0.5399 |
| 13 | Auckland Island | 0.0384 | 0.3289 | 0.3472 | 0.3231 | 0.3534 |
| 14 | Azores Canaries Madeira | 0.2562 | 0.0992 | 0.1385 | 0.1127 | 0.1190 |
| 15 | Baffin Bay - Davis Strait | 0.0920 | 0.4000 | 0.0000 | 0.3560 | 0.0000 |
| 16 | Bahamian | 0.2534 | 0.1301 | 0.1156 | 0.1264 | 0.1200 |
| 17 | Baltic Sea | 0.8491 | 0.1896 | 0.1898 | 0.1895 | 0.0902 |
| 18 | Banda Sea | 0.4112 | 0.4112 | 0.4880 | 0.4734 | 0.4563 |
| 19 | Bassian | 0.1018 | 0.3106 | 0.3046 | 0.2880 | 0.3065 |
| 20 | Beaufort-Amundsen-Viscount Melville-Queen Maud | 0.0069 | 0.0294 | 0.0000 | 0.0310 | 0.0000 |
| 21 | Beaufort Sea - continental coast and shelf | 0.0154 | 0.0886 | 0.0000 | 0.0804 | 0.0000 |
| 22 | Bermuda | 0.0884 | 0.0884 | 0.0909 | 0.1054 | 0.0872 |
| 23 | Bismarck Sea | 0.1321 | 0.3997 | 0.3935 | 0.3802 | 0.4005 |
| 24 | Black Sea | 0.5963 | 0.0943 | 0.0833 | 0.0697 | 0.1065 |
| 25 | Bonaparte Coast | 0.1399 | 0.4064 | 0.4955 | 0.4658 | 0.4534 |
| 26 | Bounty and Antipodes Island | 0.0372 | 0.3239 | 0.3525 | 0.3209 | 0.3618 |
| 27 | Bouvet Island | 0.0403 | 0.1693 | 0.3241 | 0.2800 | 0.3135 |
| 28 | Campbell Island | 0.0746 | 0.3289 | 0.3554 | 0.3159 | 0.3553 |
| 29 | Cape Howe | 0.1921 | 0.2928 | 0.2544 | 0.2680 | 0.2492 |
| 30 | Cape Verde | 0.1703 | 0.1513 | 0.1790 | 0.1423 | 0.1759 |
| 31 | Cargados Carajos/Tromelin Island | 0.1156 | 0.2912 | 0.3451 | 0.3191 | 0.3061 |
| 32 | Carolinian | 0.3027 | 0.1873 | 0.1712 | 0.2023 | 0.1627 |

| | | | | | | |
|---|---|---|---|---|---|---|
| 33 | Celtic Seas | 0.4201 | 0.3115 | 0.3670 | 0.3486 | 0.2867 |
| 34 | Central Chile | 0.1759 | 0.3623 | 0.3383 | 0.3482 | 0.3521 |
| 35 | Central Kuroshio Current | 0.4310 | 0.4358 | 0.3927 | 0.4185 | 0.4087 |
| 36 | Central New Zealand | 0.1420 | 0.3565 | 0.3430 | 0.3286 | 0.3351 |
| 37 | Central Peru | 0.1455 | 0.2887 | 0.3447 | 0.2605 | 0.3286 |
| 38 | Central Somali Coast | 0.1026 | 0.2598 | 0.2395 | 0.2633 | 0.2143 |
| 39 | Chagos | 0.1192 | 0.3260 | 0.3569 | 0.2993 | 0.3846 |
| 40 | Channels and Fjords of Southern Chile | 0.0688 | 0.3861 | 0.3580 | 0.3418 | 0.3827 |
| 41 | Chatham Island | 0.0732 | 0.3236 | 0.3261 | 0.3039 | 0.3208 |
| 42 | Chiapas-Nicaragua | 0.1734 | 0.5314 | 0.4300 | 0.4679 | 0.5185 |
| 43 | Chiloense | 0.0834 | 0.4856 | 0.4680 | 0.4651 | 0.4549 |
| 44 | Chukchi Sea | 0.0383 | 0.3632 | 0.0671 | 0.2267 | 0.0000 |
| 45 | Clipperton | 0.1474 | 0.5073 | 0.4388 | 0.5031 | 0.4900 |
| 46 | Cocos-Keeling/Christmas Island | 0.1290 | 0.3859 | 0.4050 | 0.4144 | 0.3964 |
| 47 | Cocos Islands | 0.1449 | 0.5076 | 0.5170 | 0.5073 | 0.5079 |
| 48 | Coral Sea | 0.1628 | 0.2519 | 0.2919 | 0.2667 | 0.2989 |
| 49 | Cortezian | 0.1180 | 0.3426 | 0.3424 | 0.3335 | 0.3864 |
| 50 | Crozet Islands | 0.0300 | 0.3561 | 0.3545 | 0.3374 | 0.3590 |
| 51 | Delagoa | 0.1529 | 0.2772 | 0.2334 | 0.2645 | 0.2555 |
| 52 | East Caroline Islands | 0.1175 | 0.4292 | 0.3890 | 0.4347 | 0.3888 |
| 53 | East Antarctic Enderby Land | 0.0533 | 0.0005 | 0.2073 | 0.0000 | 0.2144 |
| 54 | East Antarctic Wilkes Land | 0.0278 | 0.0084 | 0.1167 | 0.0010 | 0.1498 |
| 55 | East Antarctic Dronning Maud Land | 0.0323 | 0.0012 | 0.1449 | 0.0000 | 0.1490 |
| 56 | East China Sea | 0.7312 | 0.5451 | 0.5287 | 0.5127 | 0.5014 |
| 57 | East Greenland Shelf | 0.0985 | 0.3614 | 0.1594 | 0.2622 | 0.1413 |
| 58 | East Siberian Sea | 0.0017 | 0.1319 | 0.0000 | 0.1196 | 0.0000 |
| 59 | Easter Island | 0.0388 | 0.1959 | 0.1695 | 0.1957 | 0.1710 |
| 60 | Eastern Bering sea | 0.1348 | 0.5900 | 0.6094 | 0.6147 | 0.3546 |
| 61 | Eastern Brazil | 0.1935 | 0.0233 | 0.0271 | 0.0278 | 0.0292 |
| 62 | Eastern Caribbean | 0.2043 | 0.3217 | 0.1996 | 0.3285 | 0.1866 |
| 63 | Eastern Galapagos Islands | 0.1394 | 0.3635 | 0.3806 | 0.3386 | 0.4215 |
| 64 | Eastern India | 0.1430 | 0.4471 | 0.4882 | 0.5083 | 0.5336 |
| 65 | Eastern Philippines | 0.1541 | 0.3707 | 0.3796 | 0.3923 | 0.3913 |
| 66 | Exmouth to Broome | 0.1850 | 0.3473 | 0.3492 | 0.3574 | 0.3480 |
| 67 | Faroe Plateau | 0.1880 | 0.2715 | 0.2754 | 0.3527 | 0.1573 |
| 68 | Fernando de Naronha and Atoll das Rocas | 0.1655 | 0.1481 | 0.1759 | 0.1676 | 0.1483 |
| 69 | Fiji Islands | 0.1243 | 0.3076 | 0.3297 | 0.3122 | 0.3200 |
| 70 | Floridian | 0.3541 | 0.2605 | 0.1796 | 0.2577 | 0.2173 |
| 71 | Gilbert/Ellis Islands | 0.0473 | 0.3058 | 0.3121 | 0.3056 | 0.3122 |
| 72 | Great Australian Bight | 0.1041 | 0.2474 | 0.2174 | 0.2246 | 0.2186 |
| 73 | Guayaquil | 0.3737 | 0.3737 | 0.4469 | 0.3464 | 0.4546 |

| | | | | | | |
|---|---|---|---|---|---|---|
| 74 | Guianan | 0.1583 | 0.4361 | 0.5010 | 0.4980 | 0.3460 |
| 75 | Gulf of Aden | 0.2138 | 0.2213 | 0.1442 | 0.1133 | 0.1965 |
| 76 | Gulf of Alaska | 0.1847 | 0.5476 | 0.5039 | 0.5413 | 0.4218 |
| 77 | Gulf of Guinea Central | 0.1834 | 0.3949 | 0.4054 | 0.3792 | 0.3123 |
| 78 | Gulf of Guinea Islands | 0.1543 | 0.3520 | 0.4159 | 0.3745 | 0.4616 |
| 79 | Gulf of Guinea Upwelling | 0.1766 | 0.3451 | 0.3464 | 0.3290 | 0.3400 |
| 80 | Gulf of Guinea West | 0.1851 | 0.3676 | 0.3081 | 0.3143 | 0.3961 |
| 81 | Gulf of Maine/Bay of Fundy | 0.2555 | 0.5379 | 0.5717 | 0.5259 | 0.4889 |
| 82 | Gulf of Oman | 0.4581 | 0.2150 | 0.2379 | 0.0831 | 0.2731 |
| 83 | Gulf of Papua | 0.0891 | 0.4640 | 0.5081 | 0.4790 | 0.5125 |
| 84 | Gulf of St. Lawrence - Eastern Scotian Shelf | 0.2148 | 0.4359 | 0.5541 | 0.4328 | 0.4267 |
| 85 | Gulf of Thailan | 0.5160 | 0.5160 | 0.5347 | 0.5467 | 0.5491 |
| 86 | Gulf of Tonkin | 0.2662 | 0.5603 | 0.5166 | 0.5300 | 0.5556 |
| 87 | Halmahera | 0.1424 | 0.3969 | 0.4526 | 0.4375 | 0.4406 |
| 88 | Hawaii | 0.1539 | 0.3114 | 0.3180 | 0.3175 | 0.2978 |
| 89 | Heard and Macdonald Islands | 0.0189 | 0.3411 | 0.3524 | 0.3612 | 0.3600 |
| 90 | Houtman | 0.1941 | 0.2546 | 0.2466 | 0.2541 | 0.2364 |
| 91 | Hudson Complex | 0.0580 | 0.3013 | 0.0087 | 0.2095 | 0.0001 |
| 92 | Humboldtian | 0.1533 | 0.2826 | 0.2824 | 0.2766 | 0.2937 |
| 93 | Juan Fernandez and Desventuradas | 0.0972 | 0.3460 | 0.3307 | 0.3495 | 0.3373 |
| 94 | Kamchatka Shelf and Coast | 0.1430 | 0.5569 | 0.4477 | 0.5293 | 0.3902 |
| 95 | Kara Sea | 0.0262 | 0.2701 | 0.0011 | 0.2652 | 0.0055 |
| 96 | Kerguelen Islands | 0.0365 | 0.3729 | 0.3737 | 0.3554 | 0.3875 |
| 97 | Kermadec Island | 0.1160 | 0.2247 | 0.2252 | 0.2364 | 0.2142 |
| 98 | Lancaster Sound | 0.0472 | 0.0287 | 0.0000 | 0.0458 | 0.0000 |
| 99 | Laptev Sea | 0.0000 | 0.2569 | 0.0000 | 0.2925 | 0.0000 |
| 100 | Leeuwin | 0.1495 | 0.2590 | 0.2432 | 0.2582 | 0.2461 |
| 101 | Lesser Sunda | 0.1533 | 0.4258 | 0.4551 | 0.4738 | 0.4303 |
| 102 | Line Islands | 0.1117 | 0.3052 | 0.2761 | 0.2928 | 0.2784 |
| 103 | Lord Howe and Norfolk Islands | 0.1597 | 0.2247 | 0.2214 | 0.2272 | 0.2156 |
| 104 | Macquarie Island | 0.0234 | 0.3023 | 0.3433 | 0.3071 | 0.3407 |
| 105 | Magdalena Transition | 0.1981 | 0.3782 | 0.4161 | 0.4472 | 0.3691 |
| 106 | Malacca Strait | 0.5285 | 0.4980 | 0.5413 | 0.5516 | 0.4962 |
| 107 | Maldives | 0.1525 | 0.2336 | 0.3262 | 0.2699 | 0.3208 |
| 108 | Malvinas/Falklands | 0.0685 | 0.4532 | 0.4164 | 0.3739 | 0.4801 |
| 109 | Manning-Hawkesbury | 0.2400 | 0.2537 | 0.2481 | 0.2555 | 0.2399 |
| 110 | Mariana Islands | 0.1626 | 0.3584 | 0.3432 | 0.3746 | 0.3380 |
| 111 | Tuamotus | 0.0852 | 0.1726 | 0.1775 | 0.1671 | 0.1812 |
| 112 | Marshall Islands | 0.1065 | 0.3973 | 0.3576 | 0.3856 | 0.3545 |
| 113 | Mascarene Islands | 0.1466 | 0.2876 | 0.3117 | 0.3060 | 0.2912 |
| 114 | Mexican Tropical Pacific | 0.2025 | 0.4304 | 0.3716 | 0.4086 | 0.4048 |
| 115 | Namaqua | 0.1701 | 0.3355 | 0.3076 | 0.3102 | 0.3221 |

| | | | | | | |
|---|---|---|---|---|---|---|
| 116 | Namib | 0.1506 | 0.3507 | 0.3308 | 0.3490 | 0.3237 |
| 117 | Natal | 0.1859 | 0.2927 | 0.2490 | 0.2808 | 0.2656 |
| 118 | New Caledonia | 0.1459 | 0.2652 | 0.2900 | 0.2720 | 0.2810 |
| 119 | Nicoya | 0.1910 | 0.4508 | 0.4955 | 0.5197 | 0.4967 |
| 120 | Ningaloo | 0.1815 | 0.3047 | 0.3035 | 0.3081 | 0.2862 |
| 121 | North American Pacific Fijordland | 0.1828 | 0.5195 | 0.4615 | 0.5067 | 0.4046 |
| 122 | North Greenland | 0.0161 | 0.0287 | 0.0000 | 0.0509 | 0.0000 |
| 123 | North Patagonian Gulfs | 0.0815 | 0.6040 | 0.5640 | 0.5149 | 0.5707 |
| 124 | Northeast Sulawesi | 0.0082 | 0.4875 | 0.4912 | 0.5056 | 0.4810 |
| 125 | Northeastern Brazil | 0.1884 | 0.1015 | 0.1053 | 0.1137 | 0.0947 |
| 126 | Northeastern Honshu | 0.3092 | 0.4799 | 0.4601 | 0.4540 | 0.4223 |
| 127 | Northeastern New Zealand | 0.1439 | 0.2699 | 0.2538 | 0.2734 | 0.2416 |
| 128 | Northern and Central Red Sea | 0.2581 | 0.0164 | 0.0163 | 0.0134 | 0.0248 |
| 129 | Northern Bay of Bengal | 0.0648 | 0.2385 | 0.5231 | 0.4022 | 0.3177 |
| 130 | Northern California | 0.4516 | 0.4463 | 0.4319 | 0.4704 | 0.4158 |
| 131 | Northern Galapagos Islands | 0.1441 | 0.4640 | 0.4985 | 0.4434 | 0.4907 |
| 132 | Northern Grand Banks | 0.1619 | 0.4840 | 0.3574 | 0.4958 | 0.3898 |
| 133 | Northern Gulf of Mexico | 0.2384 | 0.2707 | 0.2302 | 0.2857 | 0.1951 |
| 134 | Northern Labrador | 0.1244 | 0.4523 | 0.0625 | 0.3905 | 0.0699 |
| 135 | Ogasawara Islands | 0.1653 | 0.3399 | 0.3111 | 0.3332 | 0.3261 |
| 136 | Oregon Washintgon Vancouver Coast and Shelf | 0.3703 | 0.5406 | 0.4868 | 0.5462 | 0.4660 |
| 137 | Oyashio Current | 0.1903 | 0.5696 | 0.4664 | 0.4803 | 0.4498 |
| 138 | Palawan/North Borneo | 0.1622 | 0.4968 | 0.4542 | 0.4750 | 0.4931 |
| 139 | Panama Bight | 0.1600 | 0.3804 | 0.5450 | 0.5263 | 0.5725 |
| 140 | Papua | 0.1001 | 0.4221 | 0.4221 | 0.4272 | 0.4248 |
| 141 | Patagonian Shelf | 0.0758 | 0.5259 | 0.4936 | 0.3936 | 0.5975 |
| 142 | Peter the First Island | 0.0169 | 0.0160 | 0.2466 | 0.0000 | 0.3105 |
| 143 | Phoenix/Tokelau/Northern Cape | 0.0716 | 0.2487 | 0.2648 | 0.2570 | 0.2484 |
| 144 | Prince Edward Islands | 0.0465 | 0.3541 | 0.3349 | 0.3226 | 0.3345 |
| 145 | Puget Trough/Georgia Basin | 1.1013 | 0.0973 | 0.0982 | 0.0977 | 0.0995 |
| 146 | Rapa-Pitcairn | 0.1041 | 0.2003 | 0.1877 | 0.1979 | 0.1913 |
| 147 | Revillagigedos | 0.1637 | 0.3931 | 0.3656 | 0.3746 | 0.3814 |
| 148 | Rio de la Plata | 0.3433 | 0.1422 | 0.1403 | 0.1432 | 0.1390 |
| 149 | Rio Grande | 0.1624 | 0.3256 | 0.2299 | 0.3183 | 0.2506 |
| 150 | Ross Sea | 0.0069 | 0.0017 | 0.0017 | 0.0000 | 0.0294 |
| 151 | Saharan Upwelling | 0.4674 | 0.1421 | 0.1892 | 0.1876 | 0.1746 |
| 152 | Samoa Islands | 0.1026 | 0.2999 | 0.3562 | 0.3397 | 0.3154 |
| 153 | Sao Pedro and Sao Paulo Islands | 0.1588 | 0.1903 | 0.1873 | 0.1841 | 0.2013 |
| 154 | Scotian Shelf | 0.2438 | 0.4584 | 0.4484 | 0.4417 | 0.5156 |
| 155 | Sea of Japan/East Sea | 0.1837 | 0.4930 | 0.3637 | 0.4476 | 0.3813 |
| 156 | Sea of Okhotsk | 0.1144 | 0.5408 | 0.3191 | 0.5139 | 0.2535 |
| 157 | Seychelles | 0.1127 | 0.2797 | 0.3017 | 0.2952 | 0.2940 |

| | | | | | | |
|---|---|---|---|---|---|---|
| 158 | Shark Bay | 0.1806 | 0.2755 | 0.2876 | 0.2781 | 0.2610 |
| 159 | Snares Island | 0.0913 | 0.3428 | 0.3424 | 0.3150 | 0.3744 |
| 160 | Society Islands | 0.1031 | 0.1706 | 0.2339 | 0.1983 | 0.2044 |
| 161 | Solomon Archipelago | 0.0937 | 0.3671 | 0.3637 | 0.3574 | 0.3637 |
| 162 | Solomon Sea | 0.1581 | 0.3626 | 0.3845 | 0.3621 | 0.3748 |
| 163 | South Australian Gulfs | 0.1508 | 0.2470 | 0.2035 | 0.2005 | 0.2264 |
| 164 | South China Sea Oceanic Island | 0.2200 | 0.4808 | 0.4346 | 0.4628 | 0.4653 |
| 165 | South European Atlantic Shelf | 0.6932 | 0.2229 | 0.2965 | 0.2846 | 0.2171 |
| 166 | South Georgia | 0.0346 | 0.3372 | 0.4031 | 0.2744 | 0.3929 |
| 167 | South India and Sri Lanka | 0.1928 | 0.4184 | 0.4073 | 0.4003 | 0.4602 |
| 168 | South Kuroshio | 0.2244 | 0.3870 | 0.3539 | 0.3837 | 0.3691 |
| 169 | South Orkney Islands | 0.0438 | 0.1402 | 0.2871 | 0.0587 | 0.3225 |
| 170 | South Sandwich Islands | 0.0477 | 0.1153 | 0.3360 | 0.0953 | 0.2868 |
| 171 | South Shetland Islands | 0.0610 | 0.1716 | 0.3206 | 0.1466 | 0.2586 |
| 172 | Southeast Madagascar | 0.1510 | 0.2997 | 0.3134 | 0.3118 | 0.3041 |
| 173 | Southeast Papua New Guinea | 0.1577 | 0.3053 | 0.3704 | 0.3004 | 0.3681 |
| 174 | Southeast Brazil | 0.2403 | 0.1997 | 0.1206 | 0.1871 | 0.1225 |
| 175 | Southern California Bright | 0.2591 | 0.3945 | 0.4115 | 0.4219 | 0.3880 |
| 176 | Southern Caribbean | 0.2203 | 0.2906 | 0.1959 | 0.2653 | 0.2201 |
| 177 | Southern China | 0.8974 | 0.5569 | 0.4838 | 0.5514 | 0.5324 |
| 178 | Southern Grand Banks | 0.2290 | 0.5193 | 0.5361 | 0.4918 | 0.4902 |
| 179 | Southern Gulf of Mexico | 0.1914 | 0.1518 | 0.1404 | 0.1462 | 0.1475 |
| 180 | South New Zealand | 0.1266 | 0.3665 | 0.3587 | 0.3207 | 0.3787 |
| 181 | Southern Red Sea | 0.2207 | 0.0370 | 0.0363 | 0.0173 | 0.0407 |
| 182 | Southern Vietnam | 0.3308 | 0.5541 | 0.4933 | 0.5511 | 0.5607 |
| 183 | Southwestern Caribbean | 0.2162 | 0.2148 | 0.1618 | 0.1728 | 0.2076 |
| 184 | St. Helena and Ascension Islands | 0.0738 | 0.1270 | 0.0988 | 0.1302 | 0.1184 |
| 185 | Sulawesi Sea/Makassar Strait | 0.1540 | 0.4311 | 0.4998 | 0.4525 | 0.5148 |
| 186 | East African Coral Coast | 0.1156 | 0.2852 | 0.3197 | 0.3101 | 0.2986 |
| 187 | Northern Monsoon Current Coast | 0.0522 | 0.2739 | 0.2601 | 0.2877 | 0.2616 |
| 188 | Bight of Sofala/Swamp Coast | 0.1309 | 0.3060 | 0.3123 | 0.3130 | 0.2930 |
| 189 | Three Kings-North Cape | 0.1636 | 0.2354 | 0.2241 | 0.2421 | 0.2249 |
| 190 | Tonga Islands | 0.1148 | 0.2655 | 0.2982 | 0.2768 | 0.2879 |
| 191 | Trindade and Martin Vaz Island | 0.0801 | 0.0288 | 0.0172 | 0.0263 | 0.0198 |
| 192 | Tristan Gough | 0.0444 | 0.3067 | 0.3202 | 0.3239 | 0.3197 |
| 193 | Marquesas | 0.0901 | 0.2183 | 0.2163 | 0.2225 | 0.2081 |
| 194 | Tweed-Moreton | 0.2222 | 0.2374 | 0.2417 | 0.2309 | 0.2508 |
| 195 | Uruguay-Buenos Aires Shelf | 0.1451 | 0.4856 | 0.4240 | 0.4580 | 0.4654 |
| 196 | Vanuatu | 0.1134 | 0.3181 | 0.3378 | 0.3258 | 0.3299 |
| 197 | Virginian | 0.4408 | 0.4426 | 0.4325 | 0.4637 | 0.3864 |
| 198 | West Caroline Islands | 0.1409 | 0.4132 | 0.4139 | 0.4273 | 0.4166 |
| 199 | Weddell Sea | 0.0160 | 0.0008 | 0.0175 | 0.0000 | 0.0226 |
| 200 | West Greenland Shelf | 0.1195 | 0.4379 | 0.1908 | 0.3746 | 0.1640 |

| | | | | | | |
|---|---|---|---|---|---|---|
| 201 | Western and Northern Madagascar | 0.1159 | 0.2877 | 0.3194 | 0.3118 | 0.3006 |
| 202 | Western Arabian Sea | 0.1883 | 0.2842 | 0.1323 | 0.0775 | 0.2306 |
| 203 | Western Bassian | 0.1217 | 0.2797 | 0.2692 | 0.2550 | 0.2765 |
| 204 | Western Caribbean | 0.2107 | 0.1813 | 0.1706 | 0.1691 | 0.2066 |
| 205 | Western Galapagos Islands | 0.1343 | 0.3413 | 0.3432 | 0.3161 | 0.3467 |
| 206 | Western India | 0.2298 | 0.2548 | 0.3089 | 0.1754 | 0.3452 |
| 207 | Yellow Sea | 0.9293 | 0.6075 | 0.5659 | 0.5557 | 0.6045 |
| 208 | Southern Java | 0.1017 | 0.4624 | 0.4656 | 0.4692 | 0.4533 |
| 209 | Western Sumatra | 0.0637 | 0.5010 | 0.4485 | 0.4572 | 0.4515 |
| 210 | Torres Strait Northern Great Barrier Reef | 0.1501 | 0.2896 | 0.3837 | 0.3000 | 0.3676 |
| 211 | Central and Southern Great Barrier Reef | 0.1869 | 0.2431 | 0.3183 | 0.2551 | 0.2980 |
| 212 | Sunda Shelf/Java Sea | 0.2201 | 0.5326 | 0.5150 | 0.5434 | 0.5348 |
| 213 | North and East Iceland | 0.1364 | 0.3119 | 0.3014 | 0.3413 | 0.2061 |
| 214 | South and West Iceland | 0.1960 | 0.3213 | 0.2834 | 0.3857 | 0.2129 |
| 215 | North Sea | 0.9756 | 0.4092 | 0.3882 | 0.4389 | 0.3571 |
| 216 | Southern Norway | 0.5805 | 0.4161 | 0.4794 | 0.4837 | 0.3107 |
| 217 | Northern Norway and Finnmark | 0.2736 | 0.4113 | 0.3883 | 0.3373 | 0.2512 |
| 218 | North and East Barents Sea | 0.0816 | 0.1974 | 0.1901 | 0.2404 | 0.1353 |
| 219 | White Sea | 0.0878 | 0.2347 | 0.0975 | 0.2334 | 0.0688 |
| 220 | Greater Antilles | 0.2214 | 0.2431 | 0.1697 | 0.1892 | 0.2185 |
| 221 | Southern Cook/Austral Islands | 0.0992 | 0.2199 | 0.2672 | 0.2470 | 0.2372 |
| 222 | Gulf of Guinea South | 0.1561 | 0.5170 | 0.5192 | 0.4972 | 0.5240 |
| 223 | Angolan | 0.1557 | 0.2669 | 0.4565 | 0.2315 | 0.3824 |
| 224 | Sahelian Upwelling | 0.2621 | 0.2824 | 0.3818 | 0.2454 | 0.3641 |
| 225 | Adriatic Sea | 0.8724 | 0.0499 | 0.0718 | 0.7386 | 0.0751 |
| 226 | Levantine Sea | 0.6236 | 0.0162 | 0.0170 | 0.0101 | 0.0214 |
| 227 | Tunisian Plateau/Gulf of Sidra | 0.2461 | 0.0167 | 0.0169 | 0.0118 | 0.0247 |
| 228 | Ionian Sea | 1.0237 | 0.0207 | 0.0267 | 0.0199 | 0.0288 |
| 229 | Aegean Sea | 1.0622 | 0.0236 | 0.0354 | 0.0268 | 0.0322 |
| 230 | Alboran Sea | 1.3245 | 0.0619 | 0.1068 | 0.1194 | 0.0877 |
| 231 | Western Mediterranean | 0.9553 | 0.0234 | 0.0479 | 0.0319 | 0.0457 |
| 232 | High Arctic Archipelago | 0.0013 | 0.0003 | 0.0001 | 0.0077 | 0.0000 |
| 233 | Humboldt Current | 0.1148 | 0.3112 | 0.2839 | 0.3048 | 0.3010 |
| 234 | Malvinas Current | 0.0708 | 0.3684 | 0.3991 | 0.3192 | 0.4024 |
| 235 | Agulhas Current | 0.1305 | 0.2635 | 0.2373 | 0.2637 | 0.2378 |
| 236 | Benguela Current | 0.1242 | 0.2470 | 0.2030 | 0.2350 | 0.2194 |
| 237 | Indian Ocean Gyre | 0.0744 | 0.2599 | 0.2532 | 0.2636 | 0.2542 |
| 238 | Leeuwin Current | 0.1522 | 0.2964 | 0.2773 | 0.2979 | 0.2727 |
| 239 | Non-gyral Southwest Pacific | 0.1540 | 0.2571 | 0.2458 | 0.2548 | 0.2440 |
| 240 | California Current | 0.2180 | 0.4597 | 0.3943 | 0.4514 | 0.4149 |
| 241 | Canary Current | 0.1831 | 0.0850 | 0.0920 | 0.0977 | 0.0917 |

| | | | | | | |
|---|---|---|---|---|---|---|
| 242 | Gulf Stream | 0.2325 | 0.3089 | 0.3155 | 0.3115 | 0.3088 |
| 243 | North Central Atlantic Gyre | 0.1711 | 0.0632 | 0.0647 | 0.0627 | 0.0645 |
| 244 | Kuroshio | 0.4222 | 0.4222 | 0.3590 | 0.3937 | 0.3807 |
| 245 | Antarctic | 0.0224 | 0.1120 | 0.2570 | 0.1236 | 0.2667 |
| 246 | Antarctic Polar Front | 0.0169 | 0.3239 | 0.3427 | 0.3295 | 0.3454 |
| 247 | Eastern Tropical Pacific | 0.1207 | 0.4073 | 0.3874 | 0.3884 | 0.3973 |
| 248 | Equatorial Atlantic | 0.1218 | 0.2036 | 0.1853 | 0.2027 | 0.1924 |
| 249 | Equatorial Pacific | 0.1015 | 0.3628 | 0.3286 | 0.3463 | 0.3329 |
| 250 | South Central Atlantic Gyre | 0.1050 | 0.1642 | 0.1443 | 0.1639 | 0.1483 |
| 251 | South Central Pacific Gyre | 0.0830 | 0.2255 | 0.2282 | 0.2309 | 0.2227 |
| 252 | Subantarctic | 0.0227 | 0.3359 | 0.3565 | 0.3392 | 0.3560 |
| 253 | Subtropical Convergence | 0.0379 | 0.3157 | 0.3230 | 0.3194 | 0.3269 |
| 254 | Indian Ocean Monsoon Gyre | 0.1269 | 0.3101 | 0.3415 | 0.3112 | 0.3194 |
| 255 | Indonesian Through-Flow | 0.1646 | 0.3888 | 0.3787 | 0.4091 | 0.3739 |
| 256 | Somali Current | 0.1741 | 0.1943 | 0.1401 | 0.0939 | 0.2172 |
| 257 | North Central Pacific Gyre | 0.1555 | 0.34739; 0 | 0.3346 | 0.3431 | 0.3326 |
| 258 | North Pacific Transitional | 0.1868 | 0.4211 | 0.3898 | 0.4204 | 0.3891 |
| 259 | Subarctic Pacific | 0.1784 | 0.4813 | 0.4128 | 0.4551 | 0.4299 |
| 260 | North Atlantic Transitional | 0.2134 | 0.2296 | 0.2379 | 0.2498 | 0.2125 |
| 261 | Subarctic Atlantic | 0.1579 | 0.3132 | 0.3044 | 0.3479 | 0.2738 |
| 262 | Arctic | 0.0134 | 0.0306 | 0.0225 | 0.0313 | 0.0156 |

**Table S2.** Predicted extra DIC exposure level and vulnerability scores of the 15 marine ecoregions around Europe (Ecological code: 17, 24, 33, 151, 165, 215, 216, 225, 226, 227, 228, 229, 230, 231, and 260) in four seasons (Dec.- Feb., Mar. – May, Jun. – Aug, and Sep. – Nov.). The ecoregions were subdivided into 9 classes based on ocean floor depth (Waller 1996) : 0 – 5m (a), 5 – 15m (b), 15 – 50 m (c), 50 – 100m (d), 100- 200m (e), 200 - 1000 m (f), 1000 – 2250 m (g), 2250 - 4500 m (h), 4500 – 11500 m (i). The risk for adverse effects from maritime shipping with underwater released CO2, as also presented in Figures S10 and S11, can be calculated by multiplying exposure with vulnerability.

| Code | Ecoregion | DIC exposure (mmol/l) | Vulnerability to algal blooms and acidification | | | |
|---|---|---|---|---|---|---|
| | | | Sep. - Nov. | Mar. - May | Jun. - Aug. | Dec. - Feb. |
| 17a | Baltic Sea | 0.5686 | 0.2103 | 0.2103 | 0.2103 | 0.1715 |
| 17b | Baltic Sea | 0.7604 | 0.2073 | 0.2074 | 0.2074 | 0.1643 |
| 17c | Baltic Sea | 1.0122 | 0.2070 | 0.2070 | 0.2070 | 0.1594 |
| 17d | Baltic Sea | 1.1768 | 0.2052 | 0.2052 | 0.2052 | 0.1335 |
| 17e | Baltic Sea | 0.9710 | 0.2069 | 0.2069 | 0.2069 | 0.1100 |
| 17f | Baltic Sea | 0.9676 | 0.2069 | 0.2070 | 0.2070 | 0.1005 |
| 24a | Black Sea | 0.3366 | 0.1420 | 0.1408 | 0.1384 | 0.1423 |
| 24b | Black Sea | 0.3075 | 0.1443 | 0.1435 | 0.1424 | 0.1445 |
| 24c | Black Sea | 0.7363 | 0.1187 | 0.1100 | 0.0948 | 0.1342 |
| 24d | Black Sea | 0.7907 | 0.0885 | 0.0929 | 0.0700 | 0.1181 |
| 24e | Black Sea | 0.7757 | 0.0868 | 0.0849 | 0.0600 | 0.1001 |
| 24f | Black Sea | 0.6776 | 0.0854 | 0.0797 | 0.0567 | 0.0984 |
| 24g | Black Sea | 0.6907 | 0.0805 | 0.0670 | 0.0540 | 0.0895 |
| 24h | Black Sea | 0.6567 | 0.0745 | 0.0701 | 0.0558 | 0.0929 |
| 33a | Celtic Seas | 0.5310 | 0.5806 | 0.5497 | 0.5852 | 0.5519 |
| 33b | Celtic Seas | 0.5159 | 0.5902 | 0.5437 | 0.5721 | 0.5553 |
| 33c | Celtic Seas | 0.6796 | 0.5692 | 0.5434 | 0.5560 | 0.5382 |
| 33d | Celtic Seas | 0.8525 | 0.4307 | 0.5191 | 0.4849 | 0.4026 |
| 33e | Celtic Seas | 0.4368 | 0.3071 | 0.4256 | 0.3503 | 0.2781 |
| 33f | Celtic Seas | 0.2003 | 0.2540 | 0.2924 | 0.3041 | 0.2147 |
| 33g | Celtic Seas | 0.1894 | 0.2450 | 0.2909 | 0.2754 | 0.2153 |
| 33h | Celtic Seas | 0.1778 | 0.2326 | 0.2737 | 0.2364 | 0.2221 |
| 151a | Saharan Upwelling | 0.6182 | 0.2965 | 0.3955 | 0.3221 | 0.2748 |
| 151b | Saharan Upwelling | 0.6963 | 0.3532 | 0.4178 | 0.3712 | 0.3243 |
| 151c | Saharan Upwelling | 0.6693 | 0.3035 | 0.4069 | 0.3486 | 0.3110 |
| 151d | Saharan Upwelling | 0.6519 | 0.2364 | 0.3533 | 0.2837 | 0.2717 |
| 151e | Saharan Upwelling | 0.8715 | 0.1889 | 0.2909 | 0.2391 | 0.2063 |
| 151f | Saharan Upwelling | 1.0273 | 0.1267 | 0.2077 | 0.1444 | 0.1290 |
| 151g | Saharan Upwelling | 0.6455 | 0.0929 | 0.1426 | 0.0549 | 0.0996 |
| 151h | Saharan Upwelling | 0.4143 | 0.0704 | 0.1194 | 0.0182 | 0.0789 |

| | | | | | | |
|---|---|---|---|---|---|---|
| 151i | Saharan Upwelling | 0.4479 | 0.0546 | 0.0970 | 0.0089 | 0.0644 |
| 165a | South European Atlantic Shelf | 0.5167 | 0.4159 | 0.5790 | 0.6029 | 0.4359 |
| 165b | South European Atlantic Shelf | 0.5528 | 0.4067 | 0.5594 | 0.5808 | 0.4171 |
| 165c | South European Atlantic Shelf | 0.7069 | 0.3921 | 0.5650 | 0.5827 | 0.4252 |
| 165d | South European Atlantic Shelf | 0.7457 | 0.3340 | 0.5364 | 0.5346 | 0.3846 |
| 165e | South European Atlantic Shelf | 0.8975 | 0.2882 | 0.4745 | 0.4121 | 0.2922 |
| 165f | South European Atlantic Shelf | 1.0662 | 0.2254 | 0.3754 | 0.2951 | 0.2062 |
| 165g | South European Atlantic Shelf | 0.9359 | 0.2207 | 0.3493 | 0.2663 | 0.1927 |
| 165h | South European Atlantic Shelf | 0.6404 | 0.1963 | 0.2895 | 0.2135 | 0.1705 |
| 165i | South European Atlantic Shelf | 0.5510 | 0.1936 | 0.2659 | 0.1948 | 0.1681 |
| 215a | North Sea | 1.1871 | 0.5481 | 0.5327 | 0.5520 | 0.4830 |
| 215b | North Sea | 1.3058 | 0.5394 | 0.4800 | 0.4938 | 0.4750 |
| 215c | North Sea | 1.4780 | 0.5190 | 0.5431 | 0.5397 | 0.4551 |
| 215d | North Sea | 0.8734 | 0.3942 | 0.4773 | 0.4599 | 0.3584 |
| 215e | North Sea | 0.6599 | 0.3369 | 0.4284 | 0.4262 | 0.3007 |
| 215f | North Sea | 0.9876 | 0.5090 | 0.3029 | 0.3125 | 0.3793 |
| 215g | North Sea | 0.2067 | 0.2643 | 0.2814 | 0.3237 | 0.2294 |
| 216a | Southern Norway | 0.6450 | 0.5174 | 0.5436 | 0.5814 | 0.1808 |
| 216b | Southern Norway | 1.4329 | 0.6003 | 0.5370 | 0.5439 | 0.3504 |
| 216c | Southern Norway | 1.4025 | 0.5841 | 0.5519 | 0.5564 | 0.3923 |
| 216d | Southern Norway | 1.3496 | 0.5894 | 0.5731 | 0.5799 | 0.4330 |
| 216e | Southern Norway | 1.4006 | 0.5909 | 0.6005 | 0.6059 | 0.4840 |
| 216f | Southern Norway | 1.2649 | 0.4884 | 0.6671 | 0.6082 | 0.5294 |
| 225a | Adriatic Sea | 0.5324 | 0.1739 | 0.2338 | 0.1805 | 0.1696 |
| 225b | Adriatic Sea | 0.4821 | 0.1472 | 0.1853 | 0.1521 | 0.1496 |
| 225c | Adriatic Sea | 0.9278 | 0.1062 | 0.1879 | 0.1010 | 0.1191 |
| 225d | Adriatic Sea | 0.9112 | 0.0394 | 0.1075 | 0.0435 | 0.0651 |
| 225e | Adriatic Sea | 0.9713 | 0.0286 | 0.0585 | 0.0257 | 0.0424 |
| 225f | Adriatic Sea | 0.7934 | 0.0340 | 0.0771 | 0.0271 | 0.0351 |
| 225g | Adriatic Sea | 0.4840 | 0.0168 | 0.0389 | 0.0131 | 0.0243 |
| 226a | Levantine Sea | 0.3497 | 0.0915 | 0.0896 | 0.0848 | 0.0958 |
| 226b | Levantine Sea | 0.5179 | 0.0961 | 0.0923 | 0.0851 | 0.0980 |
| 226c | Levantine Sea | 0.5458 | 0.0549 | 0.0532 | 0.0373 | 0.0697 |
| 226d | Levantine Sea | 0.5660 | 0.0235 | 0.0253 | 0.0173 | 0.0355 |
| 226e | Levantine Sea | 0.4854 | 0.0216 | 0.0194 | 0.0134 | 0.0251 |
| 226f | Levantine Sea | 0.4763 | 0.0251 | 0.0180 | 0.0125 | 0.0222 |

| | | | | | | |
|---|---|---|---|---|---|---|
| 226g | Levantine Sea | 0.6144 | 0.0114 | 0.0121 | 0.0068 | 0.0175 |
| 226h | Levantine Sea | 0.6968 | 0.0078 | 0.0105 | 0.0060 | 0.0160 |
| 227a | Tunisian Plateau/Gulf of Sidra | 0.1508 | 0.1106 | 0.1078 | 0.1084 | 0.1103 |
| 227b | Tunisian Plateau/Gulf of Sidra | 0.1475 | 0.0863 | 0.0726 | 0.0773 | 0.0823 |
| 227c | Tunisian Plateau/Gulf of Sidra | 0.1903 | 0.0458 | 0.0259 | 0.0245 | 0.0410 |
| 227d | Tunisian Plateau/Gulf of Sidra | 0.2116 | 0.0178 | 0.0163 | 0.0090 | 0.0291 |
| 227e | Tunisian Plateau/Gulf of Sidra | 0.2461 | 0.0118 | 0.0176 | 0.0076 | 0.0282 |
| 227f | Tunisian Plateau/Gulf of Sidra | 0.2531 | 0.0093 | 0.0122 | 0.0067 | 0.0212 |
| 227g | Tunisian Plateau/Gulf of Sidra | 0.2337 | 0.0086 | 0.0099 | 0.0060 | 0.0157 |
| 227h | Tunisian Plateau/Gulf of Sidra | 0.2936 | 0.0090 | 0.0102 | 0.0062 | 0.0159 |
| 228a | Ionian Sea | 0.7042 | 0.0862 | 0.1026 | 0.0676 | 0.1030 |
| 228b | Ionian Sea | 0.6675 | 0.0966 | 0.1258 | 0.0666 | 0.1217 |
| 228c | Ionian Sea | 0.8685 | 0.0814 | 0.1167 | 0.0594 | 0.1138 |
| 228d | Ionian Sea | 1.2857 | 0.0557 | 0.0793 | 0.0296 | 0.0709 |
| 228e | Ionian Sea | 1.2804 | 0.0524 | 0.0690 | 0.0244 | 0.0528 |
| 228f | Ionian Sea | 1.0064 | 0.0462 | 0.0578 | 0.0220 | 0.0454 |
| 228g | Ionian Sea | 0.9611 | 0.0191 | 0.0304 | 0.0156 | 0.0280 |
| 228h | Ionian Sea | 0.9978 | 0.0116 | 0.0137 | 0.0079 | 0.0196 |
| 228i | Ionian Sea | 1.3439 | 0.0118 | 0.0132 | 0.0085 | 0.0177 |
| 229a | Aegean Sea | 0.5938 | 0.0945 | 0.1169 | 0.0840 | 0.0908 |
| 229b | Aegean Sea | 0.6549 | 0.0929 | 0.1038 | 0.0728 | 0.0959 |
| 229c | Aegean Sea | 0.7644 | 0.0862 | 0.1009 | 0.0642 | 0.0959 |
| 229d | Aegean Sea | 0.9736 | 0.0573 | 0.1011 | 0.0626 | 0.0643 |
| 229e | Aegean Sea | 1.0665 | 0.0288 | 0.0666 | 0.0399 | 0.0337 |
| 229f | Aegean Sea | 1.1492 | 0.0206 | 0.0499 | 0.0307 | 0.0279 |
| 229g | Aegean Sea | 1.0416 | 0.0145 | 0.0214 | 0.0142 | 0.0237 |
| 229h | Aegean Sea | 0.9234 | 0.0103 | 0.0140 | 0.0076 | 0.0201 |
| 229i | Aegean Sea | 0.7683 | 0.0115 | 0.0132 | 0.0080 | 0.0176 |
| 230a | Alboran Sea | 0.4930 | 0.1440 | 0.2215 | 0.1360 | 0.2130 |
| 230b | Alboran Sea | 0.6102 | 0.1289 | 0.2395 | 0.1526 | 0.1969 |
| 230c | Alboran Sea | 0.6950 | 0.1313 | 0.2442 | 0.1500 | 0.1742 |
| 230d | Alboran Sea | 0.6868 | 0.1337 | 0.2520 | 0.1675 | 0.1777 |
| 230e | Alboran Sea | 0.6078 | 0.1394 | 0.2677 | 0.1951 | 0.1934 |
| 230f | Alboran Sea | 1.0687 | 0.0865 | 0.1973 | 0.1153 | 0.1162 |
| 230g | Alboran Sea | 1.5256 | 0.0454 | 0.1452 | 0.0432 | 0.0657 |
| 230h | Alboran Sea | 1.4675 | 0.0371 | 0.1289 | 0.0316 | 0.0630 |
| 231a | Western Mediterranean | 0.6245 | 0.0956 | 0.1463 | 0.0668 | 0.1216 |

| | | | | | | |
|---|---|---|---|---|---|---|
| 231b | Western Mediterranean | 0.6234 | 0.1112 | 0.1628 | 0.0728 | 0.1361 |
| 231c | Western Mediterranean | 0.7880 | 0.0757 | 0.1338 | 0.0520 | 0.1060 |
| 231d | Western Mediterranean | 1.0573 | 0.0427 | 0.0987 | 0.0348 | 0.0722 |
| 231e | Western Mediterranean | 1.1905 | 0.0344 | 0.0739 | 0.0258 | 0.0553 |
| 231f | Western Mediterranean | 1.0880 | 0.0288 | 0.0627 | 0.0223 | 0.0534 |
| 231g | Western Mediterranean | 1.0495 | 0.0232 | 0.0537 | 0.0158 | 0.0427 |
| 231h | Western Mediterranean | 0.8253 | 0.0192 | 0.0553 | 0.0123 | 0.0405 |
| 260f | Atlantic Transitional | 0.4701 | 0.0664 | 0.1215 | 0.0153 | 0.0715 |
| 260g | North Atlantic Transitional | 0.3309 | 0.2101 | 0.2475 | 0.1963 | 0.1913 |
| 260h | North Atlantic Transitional | 0.3125 | 0.2081 | 0.2755 | 0.1994 | 0.1884 |
| 260i | Atlantic Transitional | 0.2523 | 0.2081 | 0.2732 | 0.1995 | 0.1854 |
